# Differences in gut metagenomes between dairy workers and community controls: a cross-sectional study

**DOI:** 10.1101/2023.05.10.540270

**Authors:** Pauline Trinh, Marilyn C. Roberts, Peter M. Rabinowitz, Amy D. Willis

## Abstract

**Background:** As a nexus of routine antibiotic use and zoonotic pathogen presence, the livestock farming environment is a potential hotspot for the emergence of zoonotic diseases and antibiotic resistant bacteria. Livestock can further facilitate disease transmission by serving as intermediary hosts for pathogens as they undergo evolution prior to a spillover event. In light of this, we are interested in characterizing the microbiome and resistome of dairy workers, whose exposure to the livestock farming environment places them at risk for facilitating community transmission of antibiotic resistant genes and emerging zoonotic diseases.

**Results:** Using shotgun sequencing, we investigated differences in the taxonomy, diversity and gene presence of the human gut microbiome of 10 dairy farm workers and 6 community controls, supplementing these samples with additional publicly available gut metagenomes. We observed greater abundance of tetracycline resistance genes and prevalence of cephamycin resistance genes in dairy workers’ metagenomes, and lower average gene diversity. We also found evidence of commensal organism association with plasmid-mediated tetracycline resistance genes in both dairy workers and community controls (including *Faecalibacterium prausnitzii, Ligilactobacillus animalis*, and *Simiaoa sunii*). However, we did not find significant differences in the prevalence of resistance genes or virulence factors overall, nor differences in the taxonomic composition of dairy worker and community control metagenomes.

**Conclusions:** This study presents the first metagenomics analysis of United States dairy workers, providing insights into potential risks of exposure to antibiotics and pathogens in animal farming environments. Previous metagenomic studies of livestock workers in China and Europe have reported increased abundance and carriage of antibiotic resistance genes in livestock workers. While our investigation found no strong evidence for differences in the abundance or carriage of antibiotic resistance genes and virulence factors between dairy worker and community control gut metagenomes, we did observe patterns in the abundance of tetracycline resistance genes and the prevalence of cephamycin resistance genes that is consistent with previous work.

## Background

Next-generation sequencing has facilitated the study of entire microbial communities of culturable and unculturable microorganisms, revealing the profound impact that the human gut microbiome has on immune homeostasis [4, 5, 36], disease development [17, 30, 60, 91], and even resistance to pathogen invasion [1, 10, 39, 41]. The human gut microbiota is influenced by both host genetics [66, 90] and environmental factors, including diet [18, 51], geography [92], and medications [8, 53]. Recent research suggests that environmental factors outweigh host genetics in shaping the gut microbiome [29, 69]. Consequently, environments that are rich in antibiotic resistant organisms, antibiotic residues, antibiotic resistance genes (ARGs), and/or zoonotic pathogens, such as livestock farms, may pose significant risks to public health, as these environments may serve as hotspots for antibiotic resistance and zoonotic disease emergence and propagation. Studies of changes to the human microbiome and resistome in response to occupational exposure to livestock on farms may shed light on the potential risks of these environments for transmission and spread of zoonotic diseases and antibiotic resistance.

Modern farming practices and agricultural intensification have been linked to the emergence and amplification of zoonotic diseases and antimicrobial resistance (AMR), with livestock potentially serving as intermediate hosts for pathogens [38, 54]. Transmission of both zoonotic pathogens and antibiotic resistance genes can occur through direct or indirect contact at the human-animal interface, placing livestock workers and those in contact with these workers at risk of transmission and infection [31, 55]. Several shotgun metagenomic studies have looked at the effect of occupational exposure to animal agriculture on ARG carriage, finding higher prevalence of ARGs as well as evidence of transmission of ARGs from animal farming environments to workers [20, 75, 81]. While these studies highlight some potential impacts of exposure to ARG-rich animal farming environments, they either focused primarily on understanding the presence of ARGs in total community DNA without contextualizing ARGs to particular species of bacteria, or they used cultured isolates of a single bacterial species (e.g., *Escherichia coli*) to understand species-level antibiotic resistance transmission [20, 75, 81]. Furthermore, these studies did not examine virulence factor genes, which encode for functions that can cause disease and assist an organism with persisting [61]. While virulence factors have historically been associated with pathogens [61] they have also been identified on commensal or non-pathogenic genomes [32, 64], and their transmission can occur between pathogens and commensals by mobile genetic elements transmission [43, 65].

To better understand the effect of the livestock farming environment on the human gut microbiome of workers — including virulence factors, taxonomic associations of ARGs, and the role of commensal organisms in ARG transmission — we compared dairy worker and community control gut microbiomes using shotgun metagenomic sequencing. We studied differences with respect to diversity, taxonomic composition, and the carriage of virulence factor and antibiotic resistant genes. We additionally evaluated potential taxonomic affiliations of genes conferring resistance to beta-lactams (cephamycin and cephalosporins) and tetracyclines through reconstruction of their genomic context, and assessed differences in taxonomic context based on group association.

## Materials and methods

### Study participant selection

We performed metagenomic sequencing on a subset of stool samples from participants in the Healthy Dairy Worker study. The Healthy Dairy Worker study is a prospective cohort study that focuses on the effects of dairy farm exposure on the fecal and nasal microbiome, and immune and respiratory function of dairy farm workers. The study began recruitment of subjects on a rolling basis in May 2017 and involves collection of fecal and nasal samples, as well as health history data on participants at baseline enrollment, 3, 6, 12, and 24 months. Dairy workers were recruited from 3 conventional large (*>* 5, 000 animals) farms in the Yakima Valley of Eastern Washington State and community controls were recruited from surrounding communities. Recruitment of both community controls and dairy workers was done through snowball sampling where research participants assisted in identifying other potential participants. Eligibility to be a participant as a dairy worker required subjects to have been working on a dairy farm for at least 6 months. Eligibility as a community control required participants to have no prior work experience on a dairy farm in the previous 5 years, to have not lived on a dairy farm, and to have no current household member who worked on a dairy farm in the previous 5 years. Participants were consented by bilingual study staff and received an incentive payment for enrollment and subsequent sampling. Participants were asked to participate in self-reported surveys collecting information on health and work history. Sample collection and study activities were approved by the University of Washington Institutional Review Board under STUDY00000042. Study protocols have been previously described [13].

To conduct the current cross-sectional metagenomics study, we selected shotgun sequencing data of 16 fecal samples (the maximum possible with budget constraints) taken from the Healthy Dairy Worker study cohort. These samples came from 10 dairy workers and 6 community controls, all sampled at baseline enrollment. We selected the 10 dairy worker samples through simple random sampling of study subjects that met our exclusion criteria (no antibiotic use within 3 months of baseline enrollment). All dairy worker samples were selected from workers on a single farm, and all identified as white Hispanic or Latino males (both the numbers of females working on the participating dairy farms and recruitment of females into the study was low). Selection of the 6 community control samples was done using simple random sampling among community participants who had no antibiotic use within 3 months of sample collection and baseline enrollment, and who covariate-matched our dairy workers on sex and ethnicity. The unbalanced sampling of each group was designed to over-sample dairy workers, as community control samples could be supplemented with additional healthy subjects’ metagenomics data from publicly available data (i.e., The Human Microbiome Project).

Study enrollment and baseline sample collection began in 2018 for these 16 participants. The collection of study samples occurred at least one year after the Food and Drug Administration completed implementation of the Guidance for Industry (GFI) no. 213 which restricted the use of antibiotics in animal agriculture for growth promotion purposes and transitioned medically important antibiotics used in drinking water and feed from over-the-counter status to Veterinary Feed Directive (VFD) or prescription status [26, 27].

### Sampling, shotgun metagenomic library preparation and sequencing

Stool samples were self-collected by participants using a stool specimen collection kit. Participants were instructed to store stool samples in their refrigerators and to return their stool samples within 24 hours of collection to study staff. Samples were stored at *−*20*^◦^*C by field staff at a partner study site for 1-6 months before before being packaged with dry ice and transported to the University of Washington for extraction and storage at *−*20*^◦^*C. DNA extraction was performed using the MoBio DNeasy PowerLyzer PowerSoil Kit (Qiagen) following manufacturer’s protocols, and quantification of the resulting DNA was conducted using the Quant-iT PicoGreen dsDNA Assay Kit (ThermoFisher/Invitrogen). Extracted DNA samples were packaged on dry ice and transported to the Fred Hutchinson Cancer Research Center for sequencing.

Sequencing libraries were prepared from 250pg gDNA with a quarter reaction workflow using the Nextera XT Library Prep Kit (Illumina, San Diego, CA) and 12 cycles of indexing PCR. Indexed libraries were pooled by volume and post-amplification cleanup was performed with 0.8X Agencourt AMPure XP beads (Beckman Coulter, Indianapolis, IN). The library pool size distribution was validated using the Agilent High Sensitivity D5000 ScreenTape run on an Agilent 4200 TapeStation (Agilent Technologies, Inc., Santa Clara, CA). Additional library QC and cluster optimization was performed using Life Technologies-Invitrogen Qubit® 2.0 Fluorometer (Life Technologies-Invitrogen, Carlsbad, CA, USA). The resulting libraries were sequenced on the Illumina HiSeq 2500 to generate paired-end 150nt sequences for each fragment. Image analysis and base calling were performed with Illumina Real Time Analysis software v1.18.66.3, followed by demultiplexing of dual-indexed reads, removal of adapters and primers, and generation of FASTQ files with bcl2fastq Conversion Software v1.8.4 [35].

### Profiling taxonomic composition

We performed profiling of the microbial composition of the metagenomic short reads with primers and adapters removed using MetaPhlAn3 v3.0.14 [7]. MetaPhlAn3 estimates relative abundances by mapping reads to a reference database of clade-specific marker genes from ChocoPhlAn v30 (published in January 2019) [7, 72]. MetaPhlAn3 performs this read mapping against marker genes using bowtie2 v2.3.5.1 [46, 47]. Default parameters were used when running MetaPhlAn3 with an additional flag -t rel ab w read stats for outputting relative abundances with estimated number of reads mapping to each clade.

### Metagenomic assembly and processing of contigs

We conducted *de novo* assembly and processing of contigs using anvi’o v6.2 [25]. anvi’o integrates a suite of bioinformatics tools for the processing, analyzing, and visualization of metagenomics, pangenomics, and phylogenomics studies. We used the anvi’o Snakemake [44] metagenomics workflow obtained from “anvi-run-workflow” [73] with “–workflow metagenomics” to conduct our metagenomic assembly and processing of contigs. Illumina-utils [24] was used to apply the guidelines of [57] for quality filtering of reads. A median of 41.9 M (IQR: 37-47 M) reads per sample passed quality filtering. MEGAHIT v1.2.9 [49] was used to perform individual assembly of each metagenome. Further processing of the individual assemblies included generating a contigs database using anvi’o v6.2 [25], identifying open read frames using Prodigal v2.6.3 [34], predicting gene-level taxonomy using Centrifuge [42], functional annotation of genes using NCBI’s Clusters of Orthologous Groups (COGs) [76] and Pfams [23], searching for sequences using DIAMOND v0.9.14 [9], identifying single copy core genes (SCGs) using HMMER v3.3 [22] and built-in anvi’o Hidden Markov Model (HMM) profiles for bacteria and archaea, recruiting reads using bowtie2 v2.3.5.1 [46], and generating BAM files with samtools v1.10 [50]. Prediction of the approximate number of genomes in a metagenomic assembly using SCGs was done using the anvi’o script “anvi-display-contigs-stats”. Workflows using Snakemake with full parameter details can be found at the URL https://github.com/statdivlab/hdw_mgx_supplementary/.

### Metagenome annotation of virulence factors and antibiotic resistance genes

We used ABRicate v1.0.1 [71] to perform a mass screening of our *de novo* assembled gene calls for antibiotic resistance genes and virulence factor genes. ABRicate uses the Basic Local Alignment Search Tool (BLAST) [3] to annotate genes from a user-specified reference database. We used the the Virulence Factor Database v6.0 [14] and the Comprehensive Antibiotic Resistance Database (CARD) v4.0 [56] as reference databases in our search. Genes were considered present in a given metagenome if they met conservative minimum thresholds of 90% identity and 100% coverage.

Gene abundances were calculated within a metagenome by taking the mean coverage of a target ARG or VF gene divided by the sum of all mean coverages of all protein coding genes identified in a given metagenome. ARG relative abundances were further aggregated by their antibiotic classes by summing the relative abundances of genes within each antibiotic class for each metagenome. We focused our analyses to antibiotic classes that were identified by the World Health Organization (WHO) as Critically Important Antibiotics (CIA) [70].

### Comparison with the Human Microbiome Project

To supplement our community control data for comparison with our dairy worker samples, we also considered data from the Human Microbiome Project (HMP). We analyzed only HMP samples corresponding to healthy human fecal samples sampled using shotgun sequencing and samples for which there was complete participant metadata available, resulting in a total of 85 samples that were used from the HMP. These 85 samples were collected from 38 females and 47 males with an average age of 26.5 years (sd 4.7) for males and 26.0 years (sd 5.3) for females. Information on occupation was not available on participants in the HMP study, but we assume that none were dairy workers. These 85 metagenomic samples were then imported into anvi’o, where Prodigal was used to identify open reading frames. Annotation of open reading frames for virulence factors and antibiotic resistance genes from the VFDB and CARD databases was conducted using ABRicate. Taxonomic profiling of the HMP cohort was performed using MetaPhlAn3 v3.0.14 [7]. Sequencing depths for our study metagenomes ranged from 37 *−* 67 million sequenced reads per sample whereas sequencing depths for the HMP healthy cohort ranged from 21 *−* 239 million sequenced reads per sample. The average sequencing depth for the HMP healthy cohort (mean = 106M, sd = 5.9M) was higher than that of our study cohort (mean = 49M, sd = 8.7M).

### Reconstruction of genomic context of ARGs

We used our results from ABRicate to extract ARG target sequences from each metagenomic assembly. These sequences were extracted using samtools [50] and were used as “query” sequences in our genomic context reconstruction analyses. ARG query sequences were used to produce query neighborhoods that reassociated unassembled or unbinned reads that are graph-adjacent to the query sequence. To prepare our metagenomic short reads for genomic context reconstruction, we removed adapters and quality trimmed the reads using fastp [15] before removing human host reads using bbduk [12] and the masked human k-mer data [11]. Using our quality trimmed and filtered short reads and our query sequences of interest, we constructed the genomic context of each query sequence using MetaCherchant [63]. MetaCherchant uses a de Bruijn graph assembly approach to build genomic context of query sequences. We used the “environment-finder” tool in parallel and set k-mer length to 31, minimum coverage to 5, and max radius to 1000. Taxonomic annotation of sequences corresponding to graph nodes was done using kraken2 v2.1.2 [88]. Taxonomic affiliation of genes was based on kraken2 annotations of surrounding graph nodes for a particular query sequence. Identification of resistance genes located on plasmids or microbial chromosomes was conducted using the Resistance Gene Identifier (RGI) v5 [2]. The RGI integrates with the CARD database to predict AMR genes and their mutations in complete chromosome sequences, predicted genomic islands, complete plasmid sequences, and whole genome shotgun assemblies taken from National Center for Biotechnology Information (NCBI) databases. This is accomplished through prediction of open reading frames using Prodigal [34], alignment to CARD reference sequences using either BLAST [3] or DIAMOND [9], and the use of either protein homolog or protein variant models. The results from RGI’s exhaustive search are maintained and updated for each gene catalog on the CARD database.

## Results

### Study description

At the time of baseline enrollment, the dairy worker cohort had a mean of 10 years (sd 5.2) of dairy industry work. The mean age of dairy workers was lower compared to community controls (38.40 years vs. 49.50 years, t-test *p*= 0.06). We observed similar proportions of community controls who were current smokers compared to dairy workers (67% vs. 70%, Z-test *p* = 1). All community controls reported occupations as field workers in non-animal agriculture at the time of sample collection and study enrollment, which is unsurprising given the study catchment area.

### Taxonomic profiling of dairy worker and community control metagenomes

The 16 metagenomic samples were composed of 9 distinct phyla: Firmicutes, Bacteroidetes, Actinobacteria, Verrucomicrobia, Proteobacteria, Euryarchaeota, Spirochaetes, unclassified Eukaryota and Synergistetes. Of these phyla, Firmicutes, Bacteroidetes, and Actinobacteria were the 3 most abundant phyla found across all samples (Figure 1, left). The large representation of Firmicutes, Bacteroidetes, and Actinobacteria reflected similar community compositions observed in healthy subjects from the Human Microbiome Project [33]. We also note that while the majority of the phyla identified are from the domain Bacteria, we observed organisms from the domains Archaea (Euryarchaeota) and Eukaryota as well. We detected Euryarchaeota organisms in 5 dairy worker and 6 community control samples and unclassified Eukaryota organisms in low abundances in 2 dairy worker samples from our study. To examine phylum-level relative abundance differences between dairy workers and community controls, we performed a t-test of CLR-transformed read counts with pseudocounts of 1 and found no significant differences at the 5% false discovery rate level in phylum abundances.

**Figure 1:**
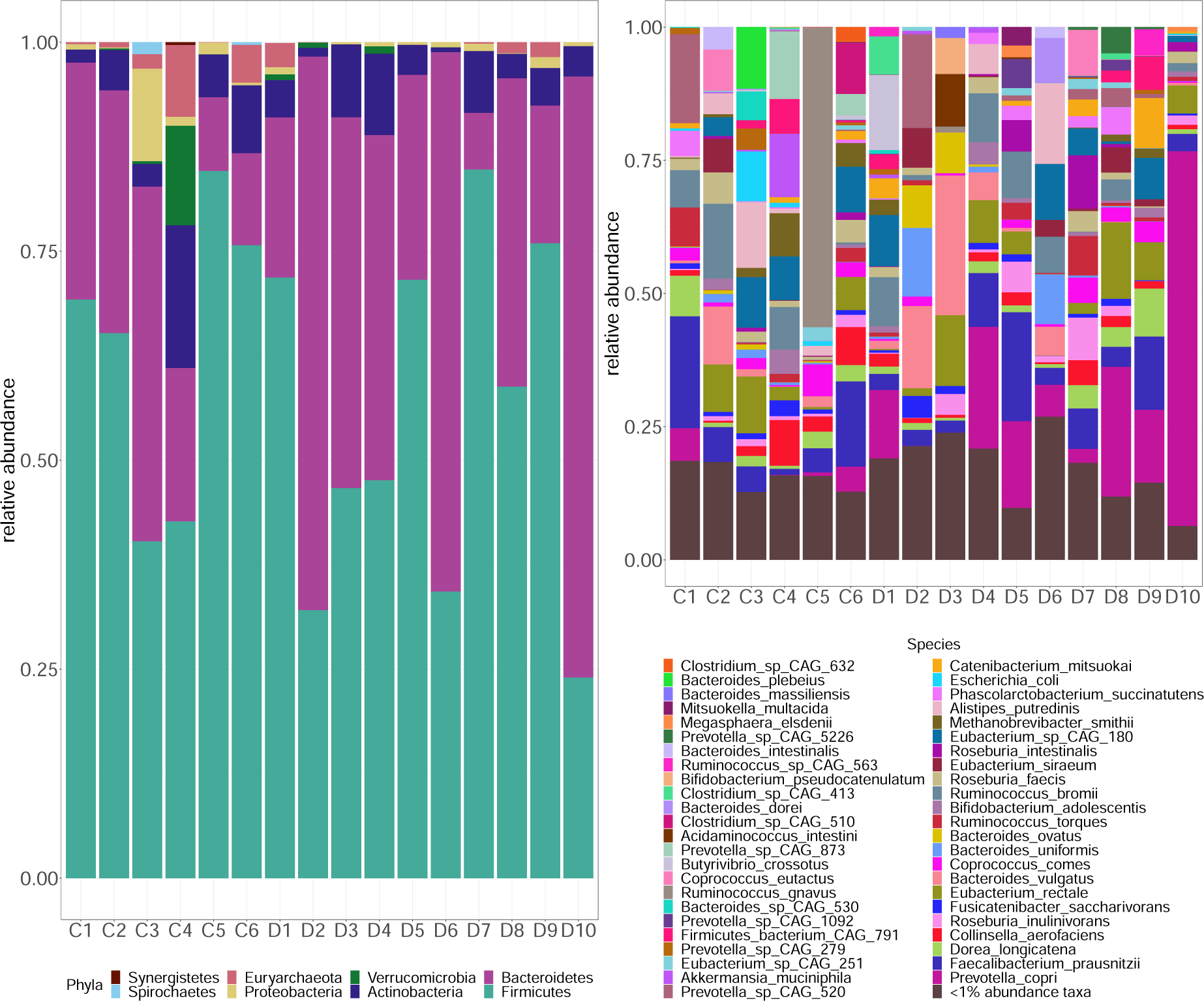
Stacked barplots of relative abundances show the most abundant phyla (left) and species (right) within each metagenome. At the phylum-level (left), Firmicutes, Bacteroidetes, and Actinobacteria are the most abundant phyla across all samples. At the species-level (right), the 5 most abundant and prevalent species across community control and dairy worker metagenomes were *F. prausnitzii, E. rectale, P. copri*, and *Eubacterium* sp. CAG-180. Species with relative abundances less than 1% were grouped together. There was insufficient evidence to suggest major differences in the taxonomic composition of dairy worker metagenomes compared to community controls.

At the species-level, we identified 272 different species across the 16 metagenomes. The most prevalent bacteria species observed were *Prevotella copri, Faecalibacterium prausnitzii, Eubacterium rectale, Ruminococcus bromii*, and *Bacteroides vulgatus* (Figure 1, right). These 5 species have been previously shown to be highly abundant organisms found in healthy human gut microflora [6, 45, 52, 78, 85]. Differential abundance testing revealed no statistically significant differences in the abundances of these 5 organisms between groups at the false discovery level of 5%. However, we did find a single organism (*Clostridium* sp. CAG 167) that was significantly less abundant in dairy worker metagenomes at the 5% FDR level (Difference in CLR-means: Δ^^^ = *−*6.93, *q* = 0.01). Abundance patterns in species with the largest magnitude test statistics (Supplementary Figure S3) showed higher abundances of *Bifidobacterium catenulatum* (Difference in CLR-means: Δ^^^ = 5.11, *q* = 0.21) and *Blautia wexlerae* (Difference in CLR-means: Δ^^^ = 1.84, *q* = 0.21) observed in dairy workers and higher abundances of *Clostridium* sp. CAG 167 (Difference in CLR-means: Δ^^^ = *−*6.93, *q* = 0.01) and *Ruminococcus callidus* (Difference in CLR-means: Δ^^^ = *−*6.03, *q* = 0.21) in community controls. We additionally conducted differential abundance testing between dairy workers and community controls with the inclusion of metagenomic data from 85 HMP healthy human subjects. A comparison of our models with and without the HMP cohort showed similar effect sizes for a given species’ CLR-transformed abundance between dairy workers and community controls in both models, but greater significance of many hypothesis tests (Supplementary Figure S2). These findings highlight the robustness of our effect sizes to the inclusion of a additional public data. Further details regarding this comparison can be found in Supplementary Figure S2.

We further investigated differences in the community structures of dairy worker and community control metagenomes by examining differences in *α−* and *β−* diversities. A comparison of the species-level *α−*diversity using Shannon diversity showed no significant difference in the *α−*diversity of dairy worker metagenomes compared to community control metagenomes (*α*^_DW_ *− α*^_CC_ = *−*0.20, *p* = 0.21). Similarly, a comparison of differences in the community composition (*β−*diversity) of dairy worker and community control metagenomes using the Bray-Curtis dissimilarity metric showed no evidence of differences in community composition between groups (Supplementary Figure S1). We additionally analyzed differences in gene-level richness (the number of unique genes) between dairy worker and community control metagenomes using geneshot [58, 59] and breakaway [86] to estimate the gene-level richness of each sample, finding significantly lower gene-level richness in dairy worker metagenomes compared to community control metagenomes (*C*^^^_DW_ *− C*^^^_CC_ *≈ −*2.0 *×* 10^5^, *p* = 0.003, [87]). To contextualize this finding, we also estimated the species richness in each metagenome using single-copy core genes [25], finding that on average there were 55 fewer species in dairy worker metagenomes compared to community control metagenomes but that this difference was not statistically significant at the 5% level (*p* = 0.31).

### Identification of virulence factor genes

Through mass screening of contigs across our 16 metagenomes using the Virulence Factor Database (VFDB) [14], we identified 37 different virulence factor genes across 4 samples (3 community control and 1 dairy worker; Supplementary Table 4, Additional File 1). We found that samples with the highest number of identified Virulence Factor Database (VFDB) genes were also those with higher sequencing depth (Figure 2, right). On average, community control samples had higher sequencing depths (mean = 2.7 *×* 10^7^, sd 3.1 *×* 10^6^) compared to dairy worker samples (mean = 2.3 *×* 10^7^, sd 4.3 *×* 10^6^) and a higher number of virulence factor genes identified than dairy workers (mean = 9.2, sd 10.1 vs. mean = 0.3, sd 0.9). Using happi [80], which accounts for unequal sequencing effort, we tested for differential enrichment of virulence factor genes between dairy worker and community control metagenomes. No virulence factor genes were significantly enriched between dairy worker and community control metagenomes at the 5% false discovery rate level (Supplementary Table 6, Additional File 1). We note that 3 community control metagenomes had higher numbers of identified virulence factor genes compared to samples of similar sequencing depth. Therefore, to contextualize our study participants among a larger set of individuals, we also considered the number of virulence factor genes identified in the Human Microbiome Project (HMP) healthy human subjects cohort. We found generally similar ranges of virulence factor genes identified between both the HMP (0 *−* 38 virulence factor genes) and HDW study (0 *−* 19 virulence factor genes) cohorts across varying sequencing depths (Figure 2, right). When we compared the male subjects from the HMP cohort to our all-male dairy worker cohort, we found that the HMP males had a range of 4-36 virulence factor genes, which was higher than the range of 0-3 virulence factor genes found in the dairy worker metagenomes. Taken together, our results do not provide strong evidence that dairy worker metagenomes contain a greater number of virulence factors than either the community controls or the HMP cohort.

**Figure 2:**
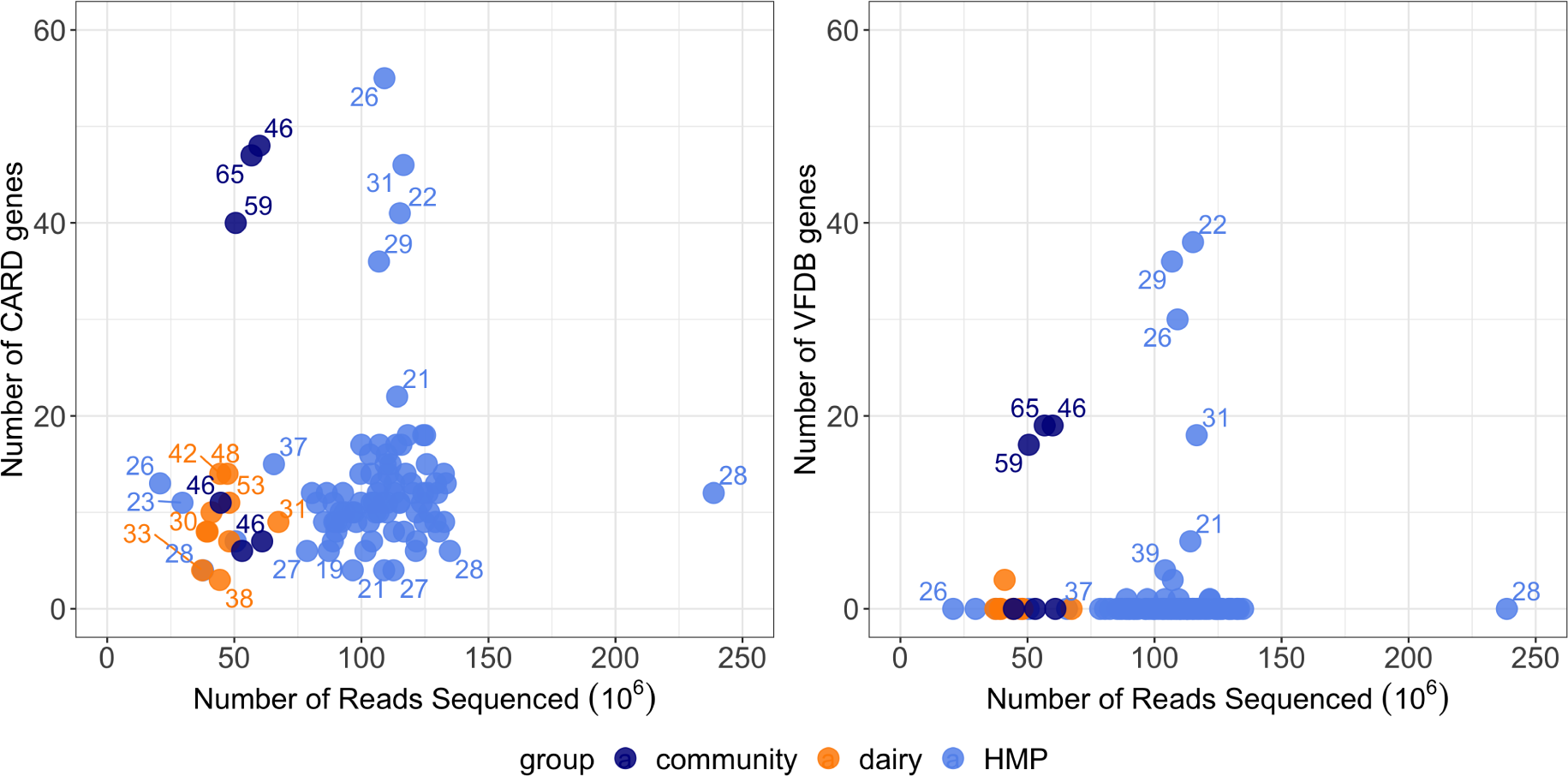
For each metagenome, we compare the sequencing depth with the number of identified CARD genes and VFDB genes. Ages (years) of each subject have been labeled. Samples with deeper sequencing had higher numbers of identified genes from the CARD and VFDB databases and higher numbers of estimated genomes. Within the community control group, 3 samples had the highest number of identified CARD genes out of all samples studied, whereas the remaining 3 community control samples within the community control group appeared to be indistinguishable from dairy workers in the number of identified CARD genes. The number of CARD and VFDB genes identified in our study cohort appeared to be similar in range to the number of CARD and VFDB genes identified in the HMP healthy human subjects cohort despite higher sequencing depths on average per sample in the HMP study cohort.

### Identification and taxonomic associations of antimicrobial resistance genes (ARGs)

Screening of the 16 metagenomes using the Comprehensive Antibiotic Resistance Database (CARD) identified 85 distinct ARGs across the 16 metagenomes, conferring resistance to at least 17 different antibiotic classes (Supplementary Figure S4; Supplementary Table 3, Additional File 1). On average, a higher number of ARGs were identified in community control metagenomes (mean = 26.5, sd 20.5) compared to dairy worker metagenomes (mean = 8.5, sd 3.7) (Figure 2, left). However, differences in the number of ARGs identified may be due, in part, to differences in sequencing depth, as metagenome samples with the highest number of ARGs identified also had higher numbers of sequenced reads (Figure 2, left). We therefore used happi to test for differences in the enrichment of ARGs between dairy worker and community control metagenomes, while accounting for differences in sequencing depth. No ARGs were differentially enriched at the false discovery level of 5%, but the following ARGs had the largest magnitude test statistics: *sat4* (happi LRT *χ*^2^ = 0.01*, q* = 0.87) a plasmid-mediated streptothricin acetyltransferase and streptothricin resistant determinant, *tet* (W) (happi LRT *χ*^2^ = 0.03*, q* = 0.87) a tetracycline resistance gene associated with both conjugative and non-conjugative DNA, and *rmtF* (happi LRT *χ*^2^ = 0.09*, q* = 0.87) an aminoglycoside resistance gene that has been found on both plasmids and chromosomes (Supplementary Table 5, Additional File 1). The 3 community metagenomes that we found to have higher numbers of virulence factor genes identified in their metagenomes also had higher numbers of ARGs identified compared to other metagenomes of similar sequencing depth. To contextualize our study cohort, we compared the number of ARGs found in our study metagenomes with the number of ARGs identified in the HMP healthy human subjects. Overall, the range of ARGs identified in our study cohort (3-48 ARGs) was similar to the range of ARGs identified in the HMP cohort (4-55 ARGs) (Figure 2, left). When we compared the male subjects from the HMP cohort to our all-male dairy worker cohort, we found that HMP males had a higher range of 4-36 ARGs compared to the range of 3-14 ARGs identified across the dairy worker metagenomes. Similar to our virulence factor genes results, these results do not provide strong evidence that dairy worker metagenomes contain greater numbers of ARGs than either community controls or the HMP cohort.

We further focused our analyses to ARGs conferring resistance to antibiotic classes considered critically important to human medicine by the World Health Organization (WHO) [70]. Across our study metagenomes, we identified 37 different ARGs conferring resistance to 8 antibiotic classes described in the WHO’s list of Critically Important Antimicrobials (CIA): aminoglycosides, fluoroquinolones, macrolides, tetracyclines, cephalosporins, cephamycins, glycopeptides, and sulfonamides (Figure 3). The most frequently occurring types of antibiotic resistance genes found across the 16 metagenomes were genes that typically confer resistance to tetracyclines (*n* = 15), aminoglycosides (*n* = 14), cephamycins (*n* = 13), and macrolides (*n* = 12) (Figure 3). Genes that commonly confer tetracycline resistance appeared to dominate the resistomes of both dairy workers and community controls with 11 distinct tetracycline resistance genes identified across 15 of our study metagenomes. We compared relative abundances of genes aggregated by antibiotic class between both groups and found that dairy workers’ metagenomes had higher mean CLR-transformed relative abundances of tetracycline (Difference in CLR-means: Δ^^^ = 0.88, *q* = 0.86), cephamycin (Difference in CLR-means: Δ^^^ = 2.74, *q* = 0.86) and macrolide (Difference in CLR-means: Δ^^^ = 0.50, *q* = 0.92) resistance genes than community controls’ metagenomes (Figure 3; Supplementary Table 8, Additional File 1); however, these differences were not significant at the 5% false discovery level. Similarly, the lower mean CLR-transformed relative abundance of aminoglycoside (Difference in CLR-means: Δ^^^ = *−*0.94, *q* = 0.86) resistance genes in dairy workers’ metagenomes compared to community controls’ metagenomes was also not significant at the 5% false discovery level.

**Figure 3:**
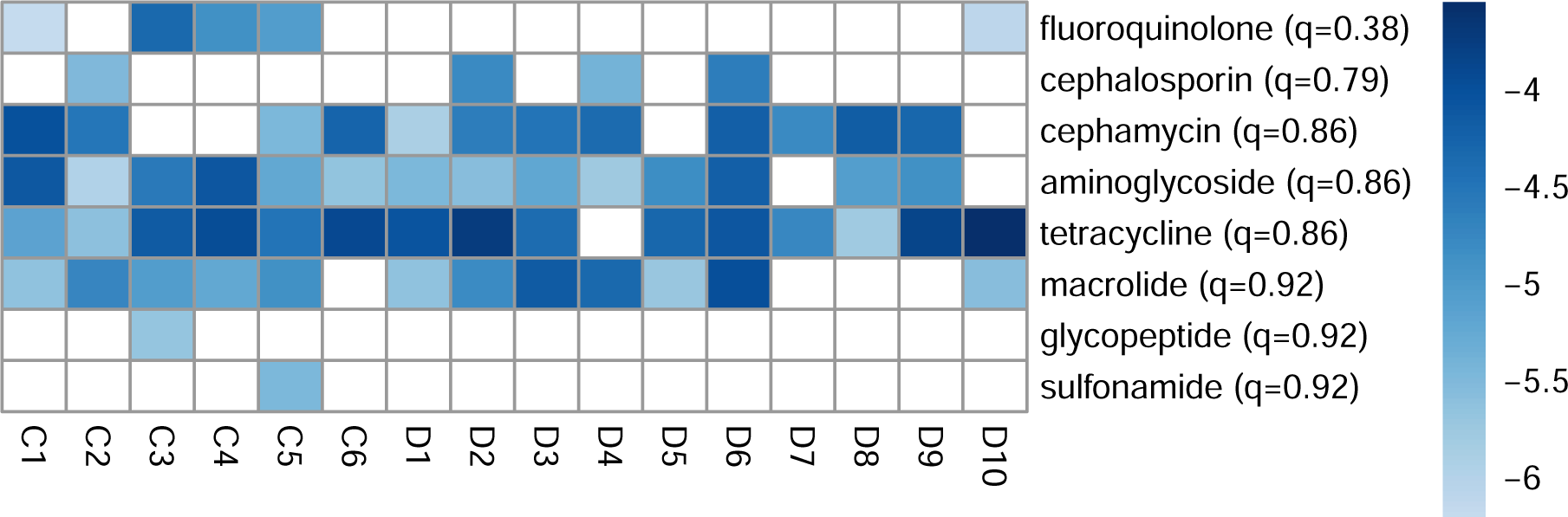
We identified ARGs from 8 antibiotic classes (rows) listed as critically important to human medicine by the WHO. log_10_ transformed relative abundances of antibiotic resistance genes grouped by these antibiotic classes are colored from lower (light blue) to higher (darker blue) relative abundances in each metagenome. Antibiotic resistance classes (rows) have been ordered by ascending q-values. White squares denote undetected antibiotic resistance genes. Visual inspection displays patterns of increased abundance of tetracycline resistance genes and macrolide resistance genes in dairy worker metagenomes. Additionally, cephamycin resistance genes had a higher occurrence in dairy workers as these genes were identified in 90% of dairy worker samples compared to 67% of community control samples.

To understand whether there were differences in taxonomic affiliation of ARGs between groups, we assessed the taxonomic context of tetracycline and beta-lactam resistance genes. We identified 6 different genes (*cblA*-1, *cfx* A2, *cfx* A3, *cfx* A4, *cfx* A5, *cfx* A6) that encode for beta-lactamases and confer resistance to beta-lactam antibiotics. Additional details on the presence of each beta-lactam resistance gene in each of our study metagenomes are found in Supplementary Table 3. These 6 beta-lactam genes have typically been identified on the chromosomes of *Bacteroides* spp. [2]. Taxonomic annotation of the genomic context of these genes in dairy worker and community control metagenomes confirmed their association with organisms from the phylum Bacteroidetes such as *Prevotella copri, Bacteroides fragilis*, and *Bacteroides uniformis*. Additionally, we observed no differences in taxonomic affiliation of these beta-lactam genes between dairy workers and community controls (Supplementary Table 7, Additional File 1).

We identified 9 *tet* genes (efflux genes: *tet* (B), *tet* (G), *tet* (40); and ribosomal genes: *tet* (M), *tet* (O), *tet* (Q), *tet* (W), *tet* (W/N/W), and *tet* (32)) that encode for efflux pumps or ribosomal protection proteins conferring resistance to tetracycline antibiotics. These genes have normally been associated with plasmids [2], which are small, extra-chromosomal DNA molecules that facilitate genetic sharing between and within species [68], but can also be found in chromosomes. Taxonomic annotation of the assembly graphs for these tetracycline resistance genes demonstrated affiliation of these genes with a variety of both commensal (e.g., *Lawsonia intracellularis, Ligilactobacillus animalis, Trueperella pyogenes, Schaalia turicensis*, and *Faecalibacterium prausnitzii*) and pathogenic (e.g., *Campylobacter* spp., *Clostridium* spp.) bacteria. Full annotations of these ARGs to affiliated bacterial organisms can be found in Supplementary Table 7, Additional File 1. Finally, while these tetracycline resistance genes were affiliated with a wide range of commensal and pathogenic bacteria, we found no differences in the taxonomic context of tetracycline resistance genes identified in community controls compared to dairy workers.

## Discussion

Using shotgun metagenomics sequencing, we investigated differences in taxonomy, diversity, and the presence of genes (especially ARGs and virulence factors) between dairy workers and community controls’ gut microbiomes. The use of shotgun metagenomics data allowed us to circumvent some of the limitations of amplicon sequencing, and enabled us to investigate abundance and presence of a variety of genes as well as their taxonomic context. To our knowledge, our study is the first shotgun metagenomics interrogation of the microbiomes and resistomes of dairy workers in the United States. While the results of our investigation revealed no statistically significant differences at the 5% level in the taxonomic composition, antibiotic resistance and virulence factor gene carriage, and relative abundances of ARGs, we observed several patterns for further investigation including greater abundance of tetracycline resistance genes and higher occurrence of cephamycin resistance genes in dairy workers’ metagenomes; evidence of commensal organism association with plasmid-mediated tetracycline resistance genes; and lower gene richness and genome diversity in dairy workers’ metagenomes.

Previous metagenomic studies of livestock workers in China and Europe have found increased abundance and carriage of antibiotic resistance genes in individuals occupationally exposed to animal farming environments, raising concerns that these environments could be hotspots for antibiotic resistance and zoonotic disease emergence [20, 75, 79, 81, 84]. Cross sectional studies of pig farmers and slaughterhouse workers in the Netherlands (*n*_workers_ = 70, *n*_controls_ = 46) [81] and China (*n*_workers_ = 4, *n*_controls_ = 5) [79] found that the resistomes of these animal workers were dominated by tetracyclines, aminoglycosides, beta-lactam and macrolide resistance genes. Another cross-sectional study of live poultry market workers in China found higher abundance of ARGs, lower Shannon diversity, and greater enrichment of beta-lactam and lincosamide resistance genes in these workers compared to controls (*n*_workers_ = 18, *n*_controls_ = 18) [84], and a longitudinal study of veterinary students with exposure to swine farms observed similar patterns of increased total abundance of ARGs and increased abundances of beta-lactam, aminoglycoside, and tetracycline resistance genes within 3 months *n* = 14 [75].

Contrary to these previous studies of livestock workers, we found no significant difference in the abundance of ARGs between dairy workers and community controls, though we did observe patterns of greater abundance of tetracycline resistant genes in dairy workers’ metagenomes that was directionally consistent with findings in these previous farm studies [20, 75, 81, 84]. In addition, we found more frequent occurrences of cephamycin (beta-lactam) resistant genes identified in the dairy worker population compared to community controls. These patterns are interesting to highlight since tetracyclines are commonly administered on dairy farms for treating gastrointestinal and respiratory diseases in dairy cows [37] and beta-lactam antibiotics such as ceftiofur are frequently used to treat metritis, a common post-partum uterine inflammatory disease [77]. It is also worth noting that the patterns observed in our study reflect the potential impacts of occupational exposure to livestock farming without the use of antibiotics for growth promotion, as the samples used in this metagenomics study were collected at least one year after the full implementation of the FDA’s GFI no. 213 policy banning the use of antibiotics for growth promotion purposes.

Our study also highlighted the potential for commensal organisms to serve as ARG reservoirs for pathogenic bacteria. By reconstructing the genomic context of each antibiotic resistance gene then taxonomically annotating this context, we were able to confirm the association of chromosome-mediated ARGs (e.g., *cblA*-1, *cfx* A2, *cfx* A3, *cfx* A4, *cfx* A5, *cfx* A6) with previously recognized carriers of these genes (e.g., *Bacteroides* spp.) [2]. With the same approach applied to primarily plasmid-mediated ARGs (e.g., *tet* (B), *tet* (G), *tet* (W/N/W), *tet* (32), *tet* (M), *tet* (O), *tet* (Q), and *tet* (W)), we found that resistance genes were associated with both commensal and pathogenic organisms. These observations suggest the potential for sharing of ARGs between commensal organisms and pathogens through conjugation. Furthermore, our results corroborate findings from a recent study that compared ARGs identified in 1,354 culture commensal strains and 45,403 pathogen strains from the human gut and found evidence of 64,188 shared ARGs that mapped to 5,931 mobile genetic elements [28]. Some of the mobile genetic elements identified [28] had also been previously identified in data from ruminant guts, soil, and other human body sites [28]. While commensal organisms may serve as ARG reservoirs for pathogenic bacteria, they may also assist in preventing pathogenic invasion through indirect (enhancement of host immune defenses) and direct (competition of nutrients and niche) mechanisms [1, 10, 39, 41]. Further research is needed to better understand the complex dynamic that commensal organisms balance in promoting both pathogen resistance and antibiotic resistance emergence.

Our results also demonstrated evidence of lower average gene richness (and some evidence of lower genome diversity) in dairy workers. Lower gene richness has been associated with increased intestinal inflammation and metabolic disorders [16, 48, 58]. A common occupational hazard facing dairy workers is inhalation of dusts and aerosols containing endotoxins or other proinflammatory substances that can result in airway inflammation and decreased pulmonary function [19, 62, 74]. Several studies have proposed a gut-lung axis linking pulmonary inflammation to intestinal inflammation based on epidemiological and clinical observations of the co-occurrence of these diseases [40, 67, 83]. There is therefore the possibility that the lower gene richness observed in dairy workers points towards increased intestinal inflammation linked to possible increased airway inflammation from exposure to aerosols and endotoxins. Further investigation to explore the possibility of increased intestinal and airway inflammation of this cohort is warranted.

Our study had several limitations. The most significant limitation was its small sample size, and therefore relatively low power to reject false null hypotheses. Corroborating our findings, especially those regarding patterns of greater tetracycline and cephamycin resistance gene in our dairy cohort, with a larger sample size is desirable. Another major limitation of our study was the comparability of the community controls with the dairy workers. The community controls in our metagenomic study occupationally identified as field workers in non-animal agriculture industries, and agricultural and dairy workers both experience occupational exposure to animal manure (e.g., as fertilizer) and antibiotics (e.g., streptomycin and oxytetracycline are commonly sprayed to control fire blight disease in Eastern Washington [21, 82]). Similar occupational exposures in the dairy workers and controls may reduce the effect sizes of group differences compared to comparisons of dairy workers and non-agricultural workers (e.g., office workers). We additionally consider the higher average age in our community controls compared to dairy workers, with 3 community controls in particular having both higher ages and the highest number of identified ARGs and virulence factors. Antibiotic resistance genes have been shown to have an age-related cumulative effect with older age groups harboring higher abundances of ARGs [89]. We were unable to determine whether the similarities between these 3 community control outliers were in part due to familial or household relatedness as this information was not collected as part of the Healthy Dairy Worker study. We did however consider the older average age as a potential driver for increased numbers of antibiotic resistance genes and virulence factors identified in these 3 community controls. We therefore provide a comparison of our dairy worker cohort to the Human Microbiome Project cohort to contextualize the dairy workers with an alternative control group. Finally, while cross-sectional studies can be advantageous for conducting cost-effective comparisons of populations, they can capture differences at a single time point. Therefore, our study cannot provide information about long-term changes to the microbiome that are induced by occupational exposure to livestock farming. We also note that shotgun metagenomics-based approaches to studying antibiotic resistance limits study to genotypic potential, and not phenotypic resistance which may not be directly correlated. Complementary future work could therefore include pairing whole genome sequencing with phenotypic resistance profiles (e.g., using culture-based approaches).

### Conclusions

In this study of occupational exposure to dairy farming, we observed no significant differences in antibiotic resistant gene or virulence factor presence in dairy workers compared to controls, but several patterns warranting further investigation, including greater abundance of tetracycline resistance genes and higher occurrence of cephamycin resistance genes in dairy workers’ metagenomes; evidence of commensal organism association with plasmid-mediated tetracycline resistance genes; and lower average gene richness in dairy workers’ metagenomes. This work demonstrates the depth and scope of utilizing shotgun metagenomics to investigate microbiomes and resistomes, and provides a foundation for further investigations into the impact of exposure to zoonotic pathogens, antibiotic resistant organisms, and ARGs on the microbiomes and resistomes of livestock workers.

## Acknowledgements

The authors would like to thank Vickie Ramirez, Jose Carmona, Lauren Frisbie, Pablo Palmandez, Theo Bammler, Pat Janssen, Doug Call, the members of COHR, Taylor Reiter, David Clausen, Sarah Teichman and members of the StatDivLab for their expert advice, support of the project, and constructive suggestions throughout the study process.

## Availability of data and materials

The data supporting the conclusions of this article along with code for reproducing our results are made available at https://github.com/statdivlab/hdw_mgx_supplementary. The sequencing data has been made available on the Sequence Read Archive under PRJNA971196.

## Abbreviations

ARG(s): Antibiotic Resistance Gene(s)
WHO: World Health Organization
HDW: Healthy Dairy Worker Study
VF: Virulence Factors
VFDB: Virulence Factor Database
CARD: Comprehensive Antibiotic Resistance Database
CIA: Critically Important Antimicrobials
LRT: Likelihood Ratio Test
CLR: Centered Log-Ratio
SCG(s): Single Copy Gene(s)
DNA: Deoxyribonucleic acid
sd: standard deviation

## Authors’ contributions

PT, PR, and AW proposed the research and wrote the manuscript. PT and AW analyzed data and created figures. All authors supported manuscript revisions, validation of results, and approved the final manuscript.

## Authors’ information

**Department of Environmental and Occupational Health Sciences, University of Washington, Seattle, WA, USA**

**Department of Biostatistics, University of Washington, Seattle, WA, USA**

Pauline Trinh

**Department of Environmental and Occupational Health Sciences, University of Washington, Seattle, WA, USA**

Marilyn C. Roberts

**Department of Environmental and Occupational Health Sciences, University of Washington, Seattle, WA, USA**

Peter M. Rabinowitz

**Department of Biostatistics, University of Washington, Seattle, WA, USA**

Amy D. Willis

## Supplementary Figures

**Figure S1:**
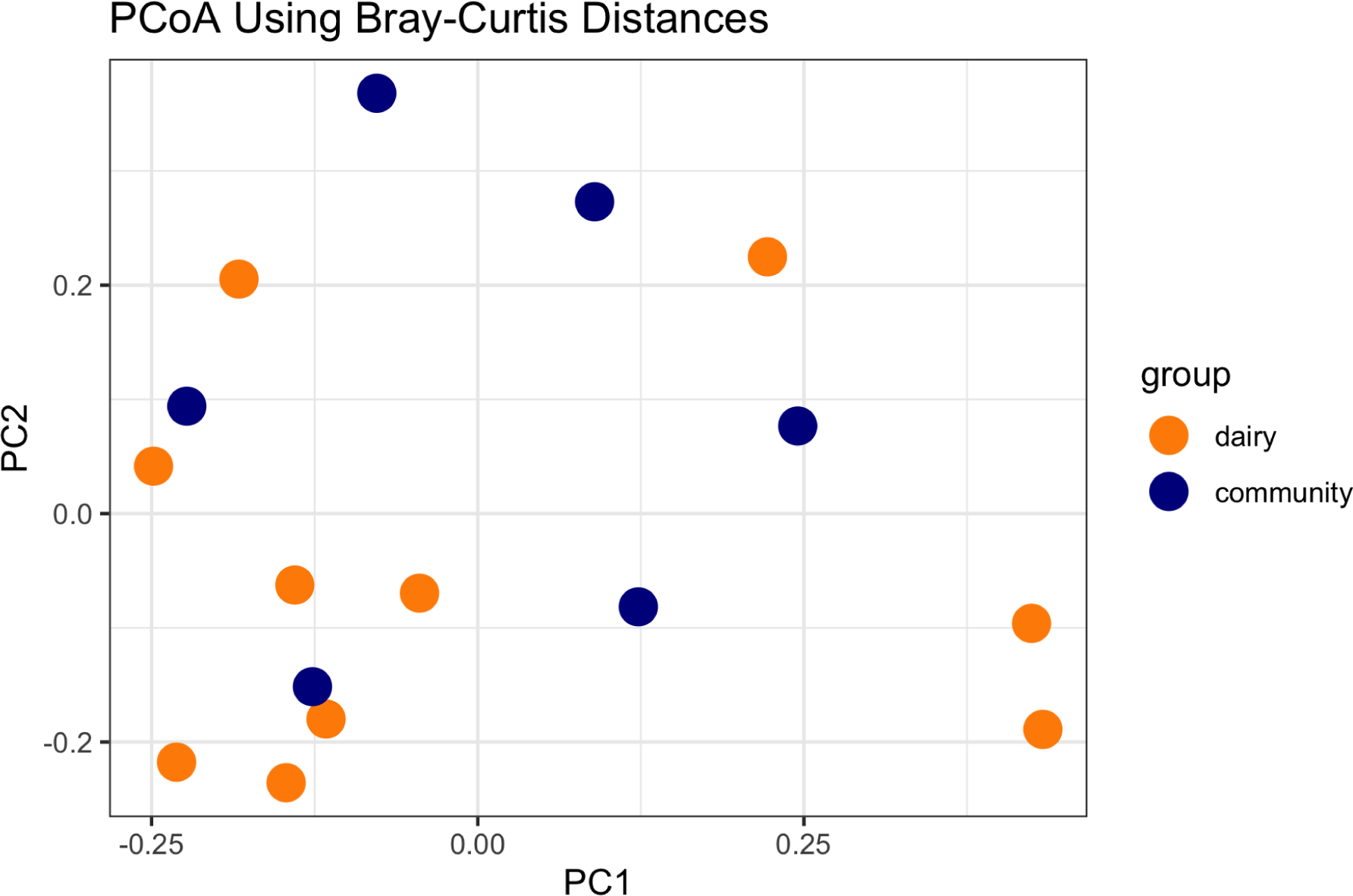
Principal coordinates analysis using Bray-Curtis distances shows similarity in microbial compositions between dairy worker and community control metagenomes.

**Figure S2:**
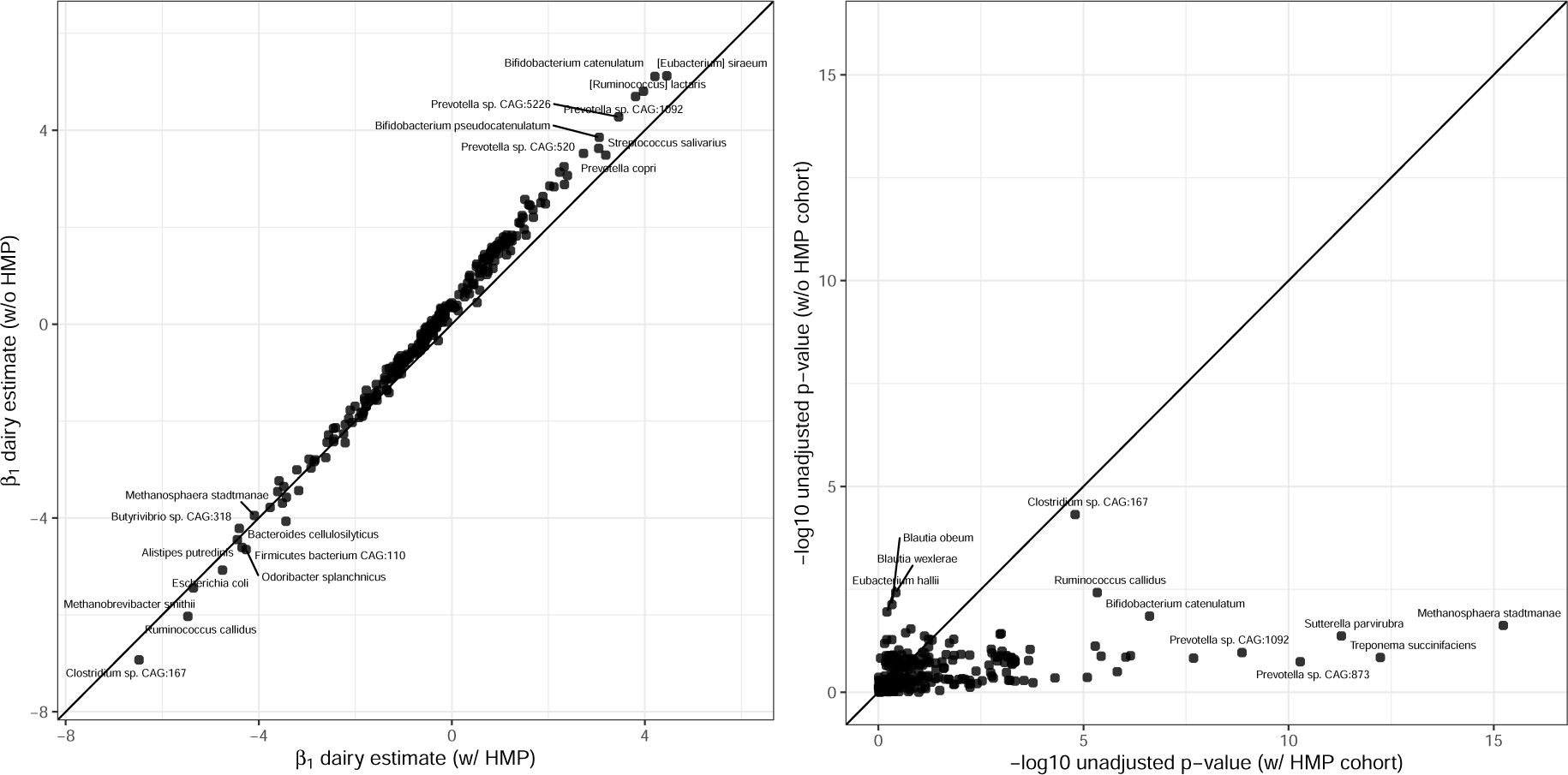
We compare parameter estimates of the difference in mean CLR-transformed abundance of species when comparing dairy workers and community controls. We contrast parameter estimates across two datasets: (1) only our study cohort, and (2) our study cohort combined with the HMP cohort. We find that coefficient estimates are robust to the inclusion of the HMP cohort (left). We also compared p-values for the null hypothesis of no difference in mean CLR abundance (right), observing that including the HMP cohort leads to more significant differences for most organisms (e.g., *M. stadtmanae, S. parvirubra*, and *T. succinifaciens*) but not all organisms (e.g., *B. wexlerae, B. obeum*, and *E. hallii*). The robustness of effect size estimates but greater significance of small p-values demonstrates that our strategy of leveraging publicly available data in conjunction with a smaller cross-sectional study can support results interpretation.

**Figure S3:**
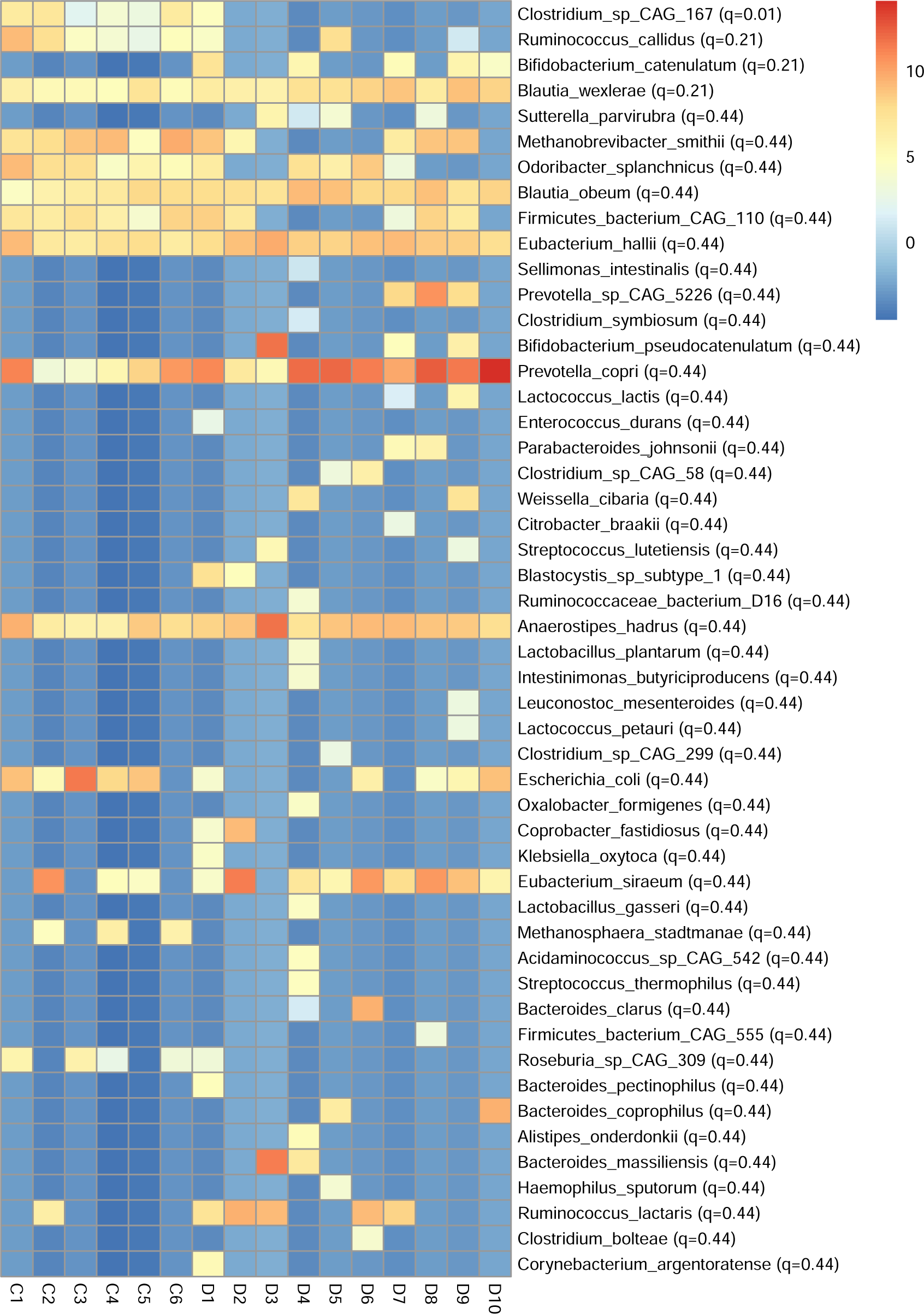
To test for differential abundance of species between dairy workers and community controls, we conducted independent t-tests of CLR-transformed relative abundances with false discovery rate correction. The heatmap displays CLR-transformed relative abundances from lower abundances (blue) to higher (red) abundances. The 50 species shown are those with the highest magnitude test statistics.

**Figure S4:**
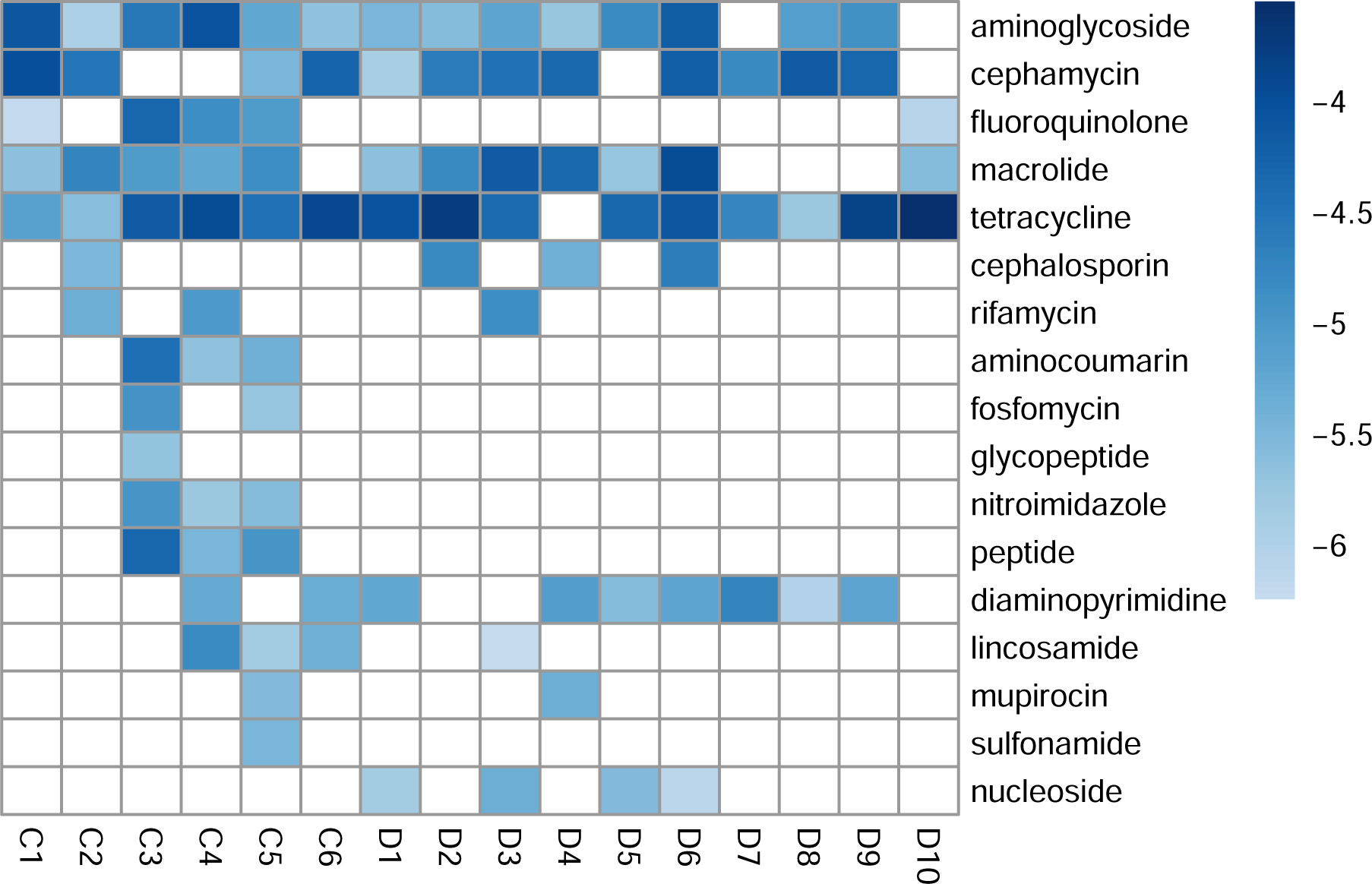
log_10_ transformed relative abundances of all antibiotic resistance genes grouped by all antibiotic classes (rows) identified across the 16 metagenomes (columns). Relative abundances are colored by magnitude, from smaller (light blue) to larger (dark blue). White cells represent that no genes were identified in the metagenome.

## References

[1] M. C. Abt and E. G. Pamer. Commensal bacteria mediated defenses against pathogens. Current Opinion in Immunology, 29:16–22, 2014. ISSN 0952-7915. doi: https://doi.org/10.1016/j.coi.2014.03.003. URL https://www.sciencedirect.com/science/article/pii/S095279151400048X.

[2] B. P. Alcock, A. R. Raphenya, T. T. Y. Lau, K. K. Tsang, M. Bouchard, A. Edalatmand, W. Huynh, A.-L. V. Nguyen, A. A. Cheng, S. Liu, S. Y. Min, A. Miroshnichenko, H.-K. Tran, R. E. Werfalli, J. A. Nasir, M. Oloni, D. J. Speicher, A. Florescu, B. Singh, M. Faltyn, A. Hernandez-Koutoucheva, A. N. Sharma, E. Bordeleau, A. C. Pawlowski, H. L. Zubyk, D. Dooley, E. Griffiths, F. Maguire, G. L. Winsor, R. G. Beiko, F. S. L. Brinkman, W. W. L. Hsiao, G. V. Domselaar, and A. G. McArthur. Card 2020: antibiotic resistome surveillance with the comprehensive antibiotic resistance database. Nucleic acids research, 48(D1):D517–D525, 01 2020. doi: 10.1093/nar/gkz935. URL https://pubmed.ncbi.nlm.nih.gov/31665441.

[3] S. Altschul, W. Gish, W. Miller, E. Myers, and D. Lipman. Basic Local Alignment Search Tool. Journal of Molecular Biology, 215(3):403–440, 1990. doi: https://doi.org/10.1016/S0022-2836(05)80360-2.

[4] M.-C. Arrieta and B. Finlay. The commensal microbiota drives immune homeostasis. Frontiers in Immunology, 3, 2012. ISSN 1664-3224. doi: 10.3389/fimmu.2012.00033. URL https://www.frontiersin.org/article/10.3389/fimmu.2012.00033.

[5] D. Artis. Epithelial-cell recognition of commensal bacteria and maintenance of immune homeostasis in the gut. Nature Reviews Immunology, 8(6):411–420, 2008. doi: 10.1038/nri2316. URL https://doi.org/10.1038/nri2316.

[6] N. T. Baxter, A. W. Schmidt, A. Venkataraman, K. S. Kim, C. Waldron, and T. M. Schmidt. Dynamics of human gut microbiota and short-chain fatty acids in response to dietary interventions with three fermentable fibers. bioRxiv, 10(1):1–13, 2018. doi: 10.1101/487900.

[7] F. Beghini, L. J. McIver, A. Blanco-Míguez, L. Dubois, F. Asnicar, S. Maharjan, A. Mailyan, P. Manghi, M. Scholz, A. M. Thomas, M. Valles-Colomer, G. Weingart, Y. Zhang, M. Zolfo, C. Huttenhower, E. A. Franzosa, and N. Segata. Integrating taxonomic, functional, and strain-level profiling of diverse microbial communities with biobakery 3. eLife, 10:1–42, 2021. doi: 10.7554/eLife.65088.

[8] M. J. Blaser. Antibiotic use and its consequences for the normal microbiome. Science (New York, N.Y.), 352(6285):544–545, 04 2016. doi: 10.1126/science.aad9358. URL https://pubmed.ncbi.nlm.nih.gov/27126037.

[9] B. Buchfink, C. Xie, and D. H. Huson. Fast and sensitive protein alignment using diamond. Nature Methods, 12(1):59–60, 2015. doi: 10.1038/nmeth.3176. URL https://doi.org/10.1038/nmeth.3176.

[10] C. G. Buffie and E. G. Pamer. Microbiota-mediated colonization resistance against intestinal pathogens. Nature Reviews Immunology, 13(11):790–801, 2013. doi: 10.1038/nri3535. URL https://doi.org/10.1038/nri3535.

[11] B. Bushnell. Masked version of hg19. URL https://zenodo.org/record/1208052#.Yq-O-i-B1Yh. Accessed: 2022-06-26.

[12] B. Bushnell. Bbtools software package, 2014. URL sourceforge.net/projects/bbmap/.

[13] J. Carmona, M. deMarcken, P. Trinh, L. Frisbie, V. Ramirez, P. Palmandez, S. Vedal, C. Sack, and P. Rabinowitz. A cross sectional study of respiratory and allergy status in dairy workers. Journal of Agromedicine, 0(0):1–8, 2023. doi: 10.1080/1059924X.2023.2171522. PMID: 36704933.

[14] L. Chen, J. Yang, J. Yu, Z. Yao, L. Sun, Y. Shen, and Q. Jin. Vfdb: a reference database for bacterial virulence factors. Nucleic Acids Research, 33(1):325–328, 01 2005. doi: 10.1093/nar/gki008. URL https://doi.org/10.1093/nar/gki008.

[15] S. Chen, Y. Zhou, Y. Chen, and J. Gu. fastp: an ultra-fast all-in-one FASTQ preprocessor. Bioinformatics, 34(17):i884–i890, 09 2018. ISSN 1367-4803. doi: 10.1093/bioinformatics/bty560. URL https://doi.org/10.1093/bioinformatics/bty560.

[16] A. Cotillard, S. P. Kennedy, L. C. Kong, E. Prifti, N. Pons, E. Le Chatelier, M. Almeida, B. Quinquis, F. Levenez, N. Galleron, S. Gougis, S. Rizkalla, J.-M. Batto, P. Renault, J. Doŕe, J.-D. Zucker, K. Cĺement, S. D. Ehrlich, H. Blottìere, M. Leclerc, C. Juste, T. de Wouters, P. Lepage, C. Fouqueray, A. Basdevant, C. Henegar, C. Godard, M. Fondacci, A. Rohia, F. Hajduch, J. Weissenbach, E. Pelletier, D. Le Paslier, J.-P. Gauchi, J.-F. Gibrat, V. Loux, W. Carŕe, E. Maguin, M. van de Guchte, A. Jamet, F. Boumezbeur, S. Layec, A. M. consortium, and A. M. consortium members. Dietary intervention impact on gut microbial gene richness. Nature, 500(7464):585–588, 2013. doi: 10.1038/nature12480. URL https://doi.org/10.1038/nature12480.

[17] A. L. Cunningham, J. W. Stephens, and D. A. Harris. Gut microbiota influence in type 2 diabetes mellitus (t2dm). Gut Pathogens, 13(1):50, 2021. doi: 10.1186/s13099-021-00446-0. URL https://doi.org/10.1186/s13099-021-00446-0.

[18] L. A. David, C. F. Maurice, R. N. Carmody, D. B. Gootenberg, J. E. Button, B. E. Wolfe, A. V. Ling, A. S. Devlin, Y. Varma, M. A. Fischbach, S. B. Biddinger, R. J. Dutton, and P. J. Turnbaugh. Diet rapidly and reproducibly alters the human gut microbiome. Nature, 505(7484): 559–563, 2014. doi: 10.1038/nature12820. URL https://doi.org/10.1038/nature12820.

[19] M. E. Davidson, J. Schaeffer, M. L. Clark, S. Magzamen, E. J. Brooks, T. J. Keefe, M. Bradford, N. Roman-Muniz, J. Mehaffy, G. Dooley, J. A. Poole, F. M. Mitloehner, S. Reed, M. B. Schenker, and S. J. Reynolds. Personal exposure of dairy workers to dust, endotoxin, muramic acid, ergosterol, and ammonia on large-scale dairies in the high plains western united states. Journal of occupational and environmental hygiene, 15(3):182–193, 03 2018. doi: 10.1080/15459624.2017.1403610. URL https://pubmed.ncbi.nlm.nih.gov/29157144.

[20] A. S. R. Duarte, T. Röder, L. Van Gompel, T. N. Petersen, R. B. Hansen, I. M. Hansen, A. Bossers, F. M. Aarestrup, J. A. Wagenaar, and T. Hald. Metagenomics-based approach to source-attribution of antimicrobial resistance determinants – identification of reservoir resistome signatures. Frontiers in Microbiology, 11, 2021. ISSN 1664-302X. doi: 10.3389/fmicb.2020.601407. URL https://www.frontiersin.org/article/10.3389/fmicb.2020.601407.

[21] B. Duffy, E. Holliger, and F. Walsh. Streptomycin use in apple orchards did not increase abundance of mobile resistance genes. FEMS Microbiology Letters, 350(2):180–189, 01 2014. ISSN 0378-1097. doi: 10.1111/1574-6968.12313. URL https://doi.org/10.1111/1574-6968.12313.

[22] S. R. Eddy. Accelerated profile hmm searches. PLOS Computational Biology, 7(10):1–16, 10 2011. doi: 10.1371/journal.pcbi.1002195. URL https://doi.org/10.1371/journal.pcbi.1002195.

[23] S. El-Gebali, J. Mistry, A. Bateman, S. R. Eddy, A. Luciani, S. C. Potter, M. Qureshi, L. J. Richardson, G. A. Salazar, A. Smart, E. L. L. Sonnhammer, L. Hirsh, L. Paladin, D. Piovesan, S. C. E. Tosatto, and R. D. Finn. The Pfam protein families database in 2019. Nucleic Acids Research, 47(D1):D427–D432, 10 2018. ISSN 0305-1048. doi: 10.1093/nar/gky995.

[24] A. M. Eren, J. H. Vineis, H. G. Morrison, and M. L. Sogin. A filtering method to generate high quality short reads using illumina paired-end technology. PLOS ONE, 8(6):1–6, 06 2013. doi: 10.1371/journal.pone.0066643. URL https://doi.org/10.1371/journal.pone.0066643.

[25] A. M. Eren, Ö. C. Esen, C. Quince, J. H. Vineis, H. G. Morrison, M. L. Sogin, and T. O. Delmont. Anvi’o: an advanced analysis and visualization platform for ‘omics data. PeerJ, 3: e1319, 2015. doi: 10.7717/peerj.1319.

[26] U. Food. Drug administration 2015. fact sheet: veterinary feed directive final rule and next steps. URL https://www.fda.gov/animal-veterinary/development-approval-process/fact-sheet-veterinary-feed-directive-final-rule-and-next-steps.

[27] U. Food, D. Administration, et al. New animal drugs and new animal drug combination products administered in or on medicated feed or drinking water of food-producing animals: recommendations for drug sponsors for voluntarily aligning product use conditions with gfi# 209. guidance for industry# 213. 2013, 2014. URL https://www.fda.gov/regulatory-information/search-fda-guidance-documents/cvm-gfi-213-new-animal-drugs-and-new-animal-drug-combination-products-administered-or-med

[28] S. C. Forster, J. Liu, N. Kumar, E. L. Gulliver, J. A. Gould, A. Escobar-Zepeda, T. Mkandawire, L. J. Pike, Y. Shao, M. D. Stares, H. P. Browne, B. A. Neville, and T. D. Lawley. Strain-level characterization of broad host range mobile genetic elements transferring antibiotic resistance from the human microbiome. Nature Communications, 13(1):1445, 2022. doi: 10.1038/s41467-022-29096-9. URL https://doi.org/10.1038/s41467-022-29096-9.

[29] R. Gacesa, A. Kurilshikov, A. Vich Vila, T. Sinha, M. A. Y. Klaassen, L. A. Bolte, S. Andreu-Śanchez, L. Chen, V. Collij, S. Hu, J. A. M. Dekens, V. C. Lenters, J. R. Björk, J. C. Swarte, M. A. Swertz, B. H. Jansen, J. Gelderloos-Arends, S. Jankipersadsing, M. Hofker, R. C. H. Vermeulen, S. Sanna, H. J. M. Harmsen, C. Wijmenga, J. Fu, A. Zhernakova, and R. K. Weersma. Environmental factors shaping the gut microbiome in a dutch population. Nature, 604(7907):732–739, 2022. doi: 10.1038/s41586-022-04567-7. URL https://doi.org/10.1038/s41586-022-04567-7.

[30] J. Geng, Q. Ni, W. Sun, L. Li, and X. Feng. The links between gut microbiota and obesity and obesity related diseases. Biomedicine and Pharmacotherapy, 147:112678, 2022. ISSN 0753-3322. doi: https://doi.org/10.1016/j.biopha.2022.112678. URL https://www.sciencedirect.com/science/article/pii/S075333222200066X.

[31] L. Grout, M. G. Baker, N. French, and S. Hales. A Review of Potential Public Health Impacts Associated With the Global Dairy Sector. GeoHealth, 4(2):1–46, 2020. ISSN 2471-1403. doi: 10.1029/2019gh000213.

[32] M. Holden, L. Crossman, A. Cerdeño-Tárraga, and J. Parkhill. Pathogenomics of non-pathogens. Nature Reviews Microbiology, 2(2):91–91, 2004. doi: 10.1038/nrmicro825. URL https://doi.org/10.1038/nrmicro825.

[33] C. Huttenhower, D. Gevers, R. Knight, S. Abubucker, J. H. Badger, A. T. Chinwalla, H. H. Creasy, A. M. Earl, M. G. FitzGerald, R. S. Fulton, M. G. Giglio, K. Hallsworth-Pepin, E. A. Lobos, R. Madupu, V. Magrini, J. C. Martin, M. Mitreva, D. M. Muzny, E. J. Sodergren, J. Versalovic, A. M. Wollam, K. C. Worley, J. R. Wortman, S. K. Young, Q. Zeng, K. M. Aagaard, O. O. Abolude, E. Allen-Vercoe, E. J. Alm, L. Alvarado, G. L. Andersen, S. Anderson, E. Appelbaum, H. M. Arachchi, G. Armitage, C. A. Arze, T. Ayvaz, C. C. Baker, L. Begg, T. Belachew, V. Bhonagiri, M. Bihan, M. J. Blaser, T. Bloom, V. Bonazzi, J. Paul Brooks, G. A. Buck, C. J. Buhay, D. A. Busam, J. L. Campbell, S. R. Canon, B. L. Cantarel, P. S. G. Chain, I.-M. A. Chen, L. Chen, S. Chhibba, K. Chu, D. M. Ciulla, J. C. Clemente, S. W. Clifton, S. Conlan, J. Crabtree, M. A. Cutting, N. J. Davidovics, C. C. Davis, T. Z. DeSantis, C. Deal, K. D. Delehaunty, F. E. Dewhirst, E. Deych, Y. Ding, D. J. Dooling, S. P. Dugan, W. Michael Dunne, A. Scott Durkin, R. C. Edgar, R. L. Erlich, C. N. Farmer, R. M. Farrell, K. Faust, M. Feldgarden, V. M. Felix, S. Fisher, A. A. Fodor, L. J. Forney, L. Foster, V. Di Francesco, J. Friedman, D. C. Friedrich, C. C. Fronick, L. L. Fulton, H. Gao, N. Garcia, G. Giannoukos, C. Giblin, M. Y. Giovanni, J. M. Goldberg, J. Goll, A. Gonzalez, A. Griggs, S. Gujja, S. Kinder Haake, B. J. Haas, H. A. Hamilton, E. L. Harris, T. A. Hepburn, B. Herter, D. E. Hoffmann, M. E. Holder, C. Howarth, K. H. Huang, S. M. Huse, J. Izard, J. K. Jansson, H. Jiang, C. Jordan, V. Joshi, J. A. Katancik, W. A. Keitel, S. T. Kelley, C. Kells, N. B. King, D. Knights, H. H. Kong, O. Koren, S. Koren, K. C. Kota, C. L. Kovar, N. C. Kyrpides, P. S. La Rosa, S. L. Lee, K. P. Lemon, N. Lennon, C. M. Lewis, L. Lewis, R. E. Ley, K. Li, K. Liolios, B. Liu, Y. Liu, C.-C. Lo, C. A. Lozupone, R. Dwayne Lunsford, T. Madden, A. A. Mahurkar, P. J. Mannon, E. R. Mardis, V. M. Markowitz, K. Mavromatis, J. M. McCorrison, D. McDonald, J. McEwen, A. L. McGuire, P. McInnes, T. Mehta, K. A. Mihindukulasuriya, J. R. Miller, P. J. Minx, I. Newsham, C. Nusbaum, M. O’Laughlin, J. Orvis, I. Pagani, K. Palaniappan, S. M. Patel, M. Pearson, J. Peterson, M. Podar, C. Pohl, K. S. Pollard, M. Pop, M. E. Priest, L. M. Proctor, X. Qin, J. Raes, J. Ravel, J. G. Reid, M. Rho, R. Rhodes, K. P. Riehle, M. C. Rivera, B. Rodriguez-Mueller, Y.-H. Rogers, M. C. Ross, C. Russ, R. K. Sanka, P. Sankar, J. Fah Sathirapongsasuti, J. A. Schloss, P. D. Schloss, T. M. Schmidt, M. Scholz, L. Schriml, A. M. Schubert, N. Segata, J. A. Segre, W. D. Shannon, R. R. Sharp, T. J. Sharpton, N. Shenoy, N. U. Sheth, G. A. Simone, I. Singh, C. S. Smillie, J. D. Sobel, D. D. Sommer, P. Spicer, G. G. Sutton, S. M. Sykes, D. G. Tabbaa, M. Thiagarajan, C. M. Tomlinson, M. Torralba, T. J. Treangen, R. M. Truty, T. A. Vishnivetskaya, J. Walker, L. Wang, Z. Wang, D. V. Ward, W. Warren, M. A. Watson, C. Wellington, K. A. Wetterstrand, J. R. White, K. Wilczek-Boney, Y. Wu, K. M. Wylie, T. Wylie, C. Yandava, L. Ye, Y. Ye, S. Yooseph, B. P. Youmans, L. Zhang, Y. Zhou, Y. Zhu, L. Zoloth, J. D. Zucker, B. W. Birren, R. A. Gibbs, S. K. Highlander, B. A. Methé, K. E. Nelson, J. F. Petrosino, G. M. Weinstock, R. K. Wilson, O. White, and T. H. M. P. Consortium. Structure, function and diversity of the healthy human microbiome. Nature, 486(7402):207–214, 2012. doi: 10.1038/nature11234. URL https://doi.org/10.1038/nature11234.

[34] D. Hyatt, G.-L. Chen, P. F. LoCascio, M. L. Land, F. W. Larimer, and L. J. Hauser. Prodigal: prokaryotic gene recognition and translation initiation site identification. BMC Bioinformatics, 11(1):119, 2010 doi: 10.1186/1471-2105-11-119. URL https://doi.org/10.1186/1471-2105-11-119.

[35] Illumina. bcl2fastq conversion software. URL http://support.illumina.com/downloads/bcl2fastq_conversion_software_184.html. Accessed: 2022-06-26.

[36] I. I. Ivanov and D. R. Littman. Modulation of immune homeostasis by commensal bacteria. Current opinion in microbiology, 14(1):106–114, 02 2011. doi: 10.1016/j.mib.2010.12.003. URL https://pubmed.ncbi.nlm.nih.gov/21215684.

[37] S. Jeamsripong, X. Li, S. Aly, Z. Su, R. Pereira, and E. Atwill. Antibiotic resistance genes and associated phenotypes in escherichia coli and enterococcus from cattle at different production stages on a dairy farm in central california. Antibiotics, 10:1042, 08 2021. doi: 10.3390/antibiotics10091042.

[38] B. A. Jones, D. Grace, R. Kock, S. Alonso, J. Rushton, M. Y. Said, D. McKeever, F. Mutua, J. Young, J. McDermott, and D. U. Pfeiffer. Zoonosis emergence linked to agricultural intensification and environmental change. Proceedings of the National Academy of Sciences of the United States of America, 110(21):8399–8404, 2013. ISSN 00278424. doi: 10.1073/pnas.1208059110.

[39] N. Kamada, S.-U. Seo, G. Y. Chen, and G. Núñez. Role of the gut microbiota in immunity and inflammatory disease. Nature Reviews Immunology, 13(5):321–335, 2013. doi: 10.1038/nri3430. URL https://doi.org/10.1038/nri3430.

[40] S. Keely, N. J. Talley, and P. M. Hansbro. Pulmonary-intestinal cross-talk in mucosal inflammatory disease. Mucosal Immunology, 5(1):7–18, 2012. doi: 10.1038/mi.2011.55. URL https://doi.org/10.1038/mi.2011.55.

[41] R. Khan, F. C. Petersen, and S. Shekhar. Commensal bacteria: An emerging player in defense against respiratory pathogens. Frontiers in Immunology, 10, 2019. ISSN 1664-3224. doi: 10.3389/fimmu.2019.01203. URL https://www.frontiersin.org/article/10.3389/fimmu.2019.01203.

[42] D. Kim, L. Song, F. P. Breitwieser, and S. L. Salzberg. Centrifuge: rapid and sensitive classification of metagenomic sequences. Genome research, 26(12):1721–1729, 12 2016. doi: 10.1101/gr.210641.116. URL https://pubmed.ncbi.nlm.nih.gov/27852649.

[43] H. Kim, M. Kim, S. Kim, Y. M. Lee, and S. C. Shin. Characterization of antimicrobial resistance genes and virulence factor genes in an arctic permafrost region revealed by metagenomics. Environmental Pollution, 294:118634, 2022. ISSN 0269-7491. doi: https://doi.org/10.1016/j.envpol.2021.118634. URL https://www.sciencedirect.com/science/article/pii/S0269749121022168.

[44] J. Köster and S. Rahmann. Snakemake—a scalable bioinformatics workflow engine. Bioinformatics, 28(19):2520–2522, 08 2012. ISSN 1367-4803. doi: 10.1093/bioinformatics/bts480. URL https://doi.org/10.1093/bioinformatics/bts480.

[45] M. Kumari, P. Singh, B. H. Nataraj, A. Kokkiligadda, H. Naithani, S. Azmal Ali, P. V. Behare, and R. Nagpal. Fostering next-generation probiotics in human gut by targeted dietary modulation: An emerging perspective. Food Research International, 150:110716, 2021. ISSN 0963-9969. doi: https://doi.org/10.1016/j.foodres.2021.110716. URL https://www.sciencedirect.com/science/article/pii/S0963996921006153.

[46] B. Langmead and S. L. Salzberg. Fast gapped-read alignment with bowtie 2. Nature Methods, 9(4):357–359, 2012. doi: 10.1038/nmeth.1923. URL https://doi.org/10.1038/nmeth.1923.

[47] B. Langmead, C. Wilks, V. Antonescu, and R. Charles. Scaling read aligners to hundreds of threads on general-purpose processors. Bioinformatics, 35(3):421–432, 07 2018. ISSN 1367-4803. doi: 10.1093/bioinformatics/bty648.

[48] E. Le Chatelier, T. Nielsen, J. Qin, E. Prifti, F. Hildebrand, G. Falony, M. Almeida, M. Arumugam, J.-M. Batto, S. Kennedy, P. Leonard, J. Li, K. Burgdorf, N. Grarup, T. Jørgensen, I. Brandslund, H. B. Nielsen, A. S. Juncker, M. Bertalan, F. Levenez, N. Pons, S. Rasmussen, S. Sunagawa, J. Tap, S. Tims, E. G. Zoetendal, S. Brunak, K. Cĺement, J. Doŕe, M. Kleerebezem, K. Kristiansen, P. Renault, T. Sicheritz-Ponten, W. M. de Vos, J.-D. Zucker, J. Raes, T. Hansen, E. Guedon, C. Delorme, S. Layec, G. Khaci, M. van de Guchte, G. Vandemeulebrouck, A. Jamet, R. Dervyn, N. Sanchez, E. Maguin, F. Haimet, Y. Winogradski, A. Cultrone, M. Leclerc, C. Juste, H. Blottìere, E. Pelletier, D. LePaslier, F. Artiguenave, T. Bruls, J. Weissenbach, K. Turner, J. Parkhill, M. Antolin, C. Manichanh, F. Casellas, N. Boruel, E. Varela, A. Torrejon, F. Guarner, G. Denariaz, M. Derrien, J. E. T. van Hylckama Vlieg, P. Veiga, R. Oozeer, J. Knol, M. Rescigno, C. Brechot, C. M’Rini, A. Mérieux, T. Yamada, P. Bork, J. Wang, S. D. Ehrlich, O. Pedersen, and M. consortium. Richness of human gut microbiome correlates with metabolic markers. Nature, 500(7464):541–546, 2013. doi: 10.1038/nature12506. URL https://doi.org/10.1038/nature12506.

[49] D. Li, C.-M. Liu, R. Luo, K. Sadakane, and T.-W. Lam. MEGAHIT: an ultra-fast single-node solution for large and complex metagenomics assembly via succinct de Bruijn graph. Bioinformatics, 31(10):1674–1676, 01 2015. ISSN 1367-4803. doi: 10.1093/bioinformatics/btv033.

[50] H. Li, B. Handsaker, A. Wysoker, T. Fennell, J. Ruan, N. Homer, G. Marth, G. Abecasis, R. Durbin, and . G. P. D. P. Subgroup. The Sequence Alignment/Map format and SAMtools. Bioinformatics, 25(16):2078–2079, 06 2009. ISSN 1367-4803. doi: 10.1093/bioinformatics/btp352. URL https://doi.org/10.1093/bioinformatics/btp352.

[51] H. Liu, M. Han, S. C. Li, G. Tan, S. Sun, Z. Hu, P. Yang, R. Wang, Y. Liu, F. Chen, J. Peng, H. Peng, H. Song, Y. Xia, L. Chu, Q. Zhou, F. Guan, J. Wu, D. Bu, and K. Ning. Resilience of human gut microbial communities for the long stay with multiple dietary shifts. Gut, 68 (12):2254–2255, 2019. ISSN 0017-5749. doi: 10.1136/gutjnl-2018-317298.

[52] D. Machado, J. C. Barbosa, M. Domingos, D. Almeida, J. C. Andrade, A. C. Freitas, and A. M. Gomes. Revealing antimicrobial resistance profile of the novel probiotic candidate faecalibacterium prausnitzii dsm 17677. International Journal of Food Microbiology, 363: 109501, 2022. ISSN 0168-1605. doi: https://doi.org/10.1016/j.ijfoodmicro.2021.109501. URL https://www.sciencedirect.com/science/article/pii/S0168160521004608.

[53] L. Maier, M. Pruteanu, M. Kuhn, G. Zeller, A. Telzerow, E. E. Anderson, A. R. Brochado, K. C. Fernandez, H. Dose, H. Mori, K. R. Patil, P. Bork, and A. Typas. Extensive impact of non-antibiotic drugs on human gut bacteria. Nature, 555(7698):623–628, 2018. doi: 10.1038/nature25979. URL https://doi.org/10.1038/nature25979.

[54] C. Manyi-Loh, S. Mamphweli, E. Meyer, and A. Okoh. Antibiotic use in agriculture and its consequential resistance in environmental sources: Potential public health implications. Molecules, 23(4), 2018. ISSN 1420-3049. doi: 10.3390/molecules23040795. URL https://www.mdpi.com/1420-3049/23/4/795.

[55] B. M. Marshall and S. B. Levy. Food animals and antimicrobials: Impacts on human health. Clinical Microbiology Reviews, 24(4):718–733, 2011. ISSN 08938512. doi: 10.1128/CMR.00002-11.

[56] A. G. McArthur, N. Waglechner, F. Nizam, A. Yan, M. A. Azad, A. J. Baylay, K. Bhullar, M. J. Canova, G. De Pascale, L. Ejim, L. Kalan, A. M. King, K. Koteva, M. Morar, M. R. Mulvey, J. S. O’Brien, A. C. Pawlowski, L. J. V. Piddock, P. Spanogiannopoulos, A. D. Sutherland, I. Tang, P. L. Taylor, M. Thaker, W. Wang, M. Yan, T. Yu, and G. D. Wright. The Comprehensive Antibiotic Resistance Database. Antimicrobial Agents and Chemotherapy, 57(7):3348–3357, 2013. ISSN 0066-4804. doi: 10.1128/aac.00419-13.

[57] A. Minoche, J. C. Dohm, and H. Himmelbauer. Evaluation of genomic high-throughput sequencing data generated on illumina hiseq and genome analyzer systems. Genome biology, 12(11):R112–R112, 11 2011. doi: 10.1186/gb-2011-12-11-r112. URL https://pubmed.ncbi.nlm.nih.gov/22067484.

[58] S. S. Minot and A. D. Willis. Clustering co-abundant genes identifies components of the gut microbiome that are reproducibly associated with colorectal cancer and inflammatory bowel disease. Microbiome, 7(1):110, 2019. doi: 10.1186/s40168-019-0722-6. URL https://doi.org/10.1186/s40168-019-0722-6.

[59] S. S. Minot, K. C. Barry, C. Kasman, J. L. Golob, and A. D. Willis. Geneshot: Gene-Level Metagenomics Identifies Genome Islands Associated With Immunotherapy Response. Genome Biology, 22(1):1–10, 2021. doi: 10.1186/s13059-021-02355-6.

[60] Y. Nakanishi, T. Sato, and T. Ohteki. Commensal gram-positive bacteria initiates colitis by inducing monocyte/macrophage mobilization. Mucosal Immunology, 8(1):152–160, 2015. doi: 10.1038/mi.2014.53. URL https://doi.org/10.1038/mi.2014.53.

[61] C. Niu, D. Yu, Y. Wang, H. Ren, Y. Jin, W. Zhou, B. Li, Y. Cheng, J. Yue, Z. Gao, and L. Liang. Common and pathogen-specific virulence factors are different in function and structure. Virulence, 4(6):473–482, 2013. doi: 10.4161/viru.25730. PMID: 23863604.

[62] M. W. Nonnenmann, D. Gimeno Ruiz de Porras, J. Levin, D. Douphrate, V. Boggaram, J. Schaffer, M. Gallagher, M. Hornick, and S. Reynolds. Pulmonary function and airway inflammation among dairy parlor workers after exposure to inhalable aerosols. American journal of industrial medicine, 60(3):255–263, 03 2017. doi: 10.1002/ajim.22680. URL https://pubmed.ncbi.nlm.nih.gov/28195657.

[63] E. I. Olekhnovich, A. T. Vasilyev, V. I. Ulyantsev, E. S. Kostryukova, and A. V. Tyakht. MetaCherchant: Analyzing genomic context of antibiotic resistance genes in gut microbiota. Bioinformatics, 34(3):434–444, 2018. ISSN 14602059. doi: 10.1093/bioinformatics/btx681.

[64] M. J. Pallen and B. W. Wren. Bacterial pathogenomics. Nature, 449(7164):835–842, 2007. doi: 10.1038/nature06248. URL https://doi.org/10.1038/nature06248.

[65] Y. Pan, J. Zeng, L. Li, J. Yang, Z. Tang, W. Xiong, Y. Li, S. Chen, Z. Zeng, and J. A. Gilbert. Coexistence of antibiotic resistance genes and virulence factors deciphered by largescale complete genome analysis. mSystems, 5(3):e00821–19, 2020. doi: 10.1128/mSystems.00821-19.

[66] Y. Qin, A. S. Havulinna, Y. Liu, P. Jousilahti, S. C. Ritchie, A. Tokolyi, J. G. Sanders, L. Valsta, M. Brożyńska, Q. Zhu, A. Tripathi, Y. Vázquez-Baeza, R. Loomba, S. Cheng, M. Jain, T. Niiranen, L. Lahti, R. Knight, V. Salomaa, M. Inouye, and G. Méric. Combined effects of host genetics and diet on human gut microbiota and incident disease in a single population cohort. Nature Genetics, 54(2):134–142, 2022. doi: 10.1038/s41588-021-00991-z. URL https://doi.org/10.1038/s41588-021-00991-z.

[67] A. L. Raftery, E. Tsantikos, N. L. Harris, and M. L. Hibbs. Links between inflammatory bowel disease and chronic obstructive pulmonary disease. Frontiers in immunology, 11:2144–2144, 09 2020. doi: 10.3389/fimmu.2020.02144. URL https://pubmed.ncbi.nlm.nih.gov/33042125.

[68] J. Rodŕıguez-Beltŕan, J. DelaFuente, R. Léon-Sampedro, R. C. MacLean, and Á. San Milĺan. Beyond horizontal gene transfer: the role of plasmids in bacterial evolution. Nature Reviews Microbiology, 19(6):347–359, 2021. doi: 10.1038/s41579-020-00497-1. URL https://doi.org/10.1038/s41579-020-00497-1.

[69] D. Rothschild, O. Weissbrod, E. Barkan, A. Kurilshikov, T. Korem, D. Zeevi, P. I. Costea, A. Godneva, I. N. Kalka, N. Bar, S. Shilo, D. Lador, A. V. Vila, N. Zmora, M. Pevsner-Fischer, D. Israeli, N. Kosower, G. Malka, B. C. Wolf, T. Avnit-Sagi, M. Lotan-Pompan, A. Weinberger, Z. Halpern, S. Carmi, J. Fu, C. Wijmenga, A. Zhernakova, E. Elinav, and E. Segal. Environment dominates over host genetics in shaping human gut microbiota. Nature, 555(7695): 210–215, 2018. doi: 10.1038/nature25973. URL https://doi.org/10.1038/nature25973.

[70] H. M. Scott, G. Acuff, G. Bergeron, M. W. Bourassa, J. Gill, D. W. Graham, L. H. Kahn, P. S. Morley, M. J. Salois, S. Simjee, R. S. Singer, T. C. Smith, C. Storrs, and T. E. Wittum. Critically important antibiotics: criteria and approaches for measuring and reducing their use in food animal agriculture. Annals of the New York Academy of Sciences, 1441(1):8–16, 2019. doi: https://doi.org/10.1111/nyas.14058. URL https://nyaspubs.onlinelibrary.wiley.com/doi/abs/10.1111/nyas.14058.

[71] T. Seemann. Abricate: mass screening of contigs for antiobiotic resistance genes. 2016. URL https://github.com/tseemann/abricate.

[72] N. Segata, L. Waldron, A. Ballarini, V. Narasimhan, O. Jousson, and C. Huttenhower. Metagenomic microbial community profiling using unique clade-specific marker genes. Nature Methods, 9(8):811–814, 2012. ISSN 15487091. doi: 10.1038/nmeth.2066.

[73] A. Shaiber, A. D. Willis, T. O. Delmont, S. Roux, L. X. Chen, A. C. Schmid, M. Yousef, A. R. Watson, K. Lolans, Ö. C. Esen, S. T. M. Lee, N. Downey, H. G. Morrison, F. E. Dewhirst, J. L. Mark Welch, and A. M. Eren. Functional and genetic markers of niche partitioning among enigmatic members of the human oral microbiome. Genome Biology, page 21:292, 2020. ISSN 26928205. doi: 10.1101/2020.04.29.069278.

[74] S. Stoleski, J. Minov, J. Karadzinska-Bislimovska, D. Mijakoski, A. Atanasovska, and D. Bislimovska. Asthma and chronic obstructive pulmonary disease associated with occupational exposure in dairy farmers - importance of job exposure matrices. Open access Macedonian journal of medical sciences, 7(14):2350–2359, 07 2019. doi: 10.3889/oamjms.2019.630. URL https://pubmed.ncbi.nlm.nih.gov/31592062.

[75] J. Sun, X.-P. Liao, A. W. D’Souza, M. Boolchandani, S.-H. Li, K. Cheng, J. Luis Martínez, L. Li, Y.-J. Feng, L.-X. Fang, T. Huang, J. Xia, Y. Yu, Y.-F. Zhou, Y.-X. Sun, X.-B. Deng, Z.-L. Zeng, H.-X. Jiang, B.-H. Fang, Y.-Z. Tang, X.-L. Lian, R.-M. Zhang, Z.-W. Fang, Q.-L. Yan, G. Dantas, and Y.-H. Liu. Environmental remodeling of human gut microbiota and antibiotic resistome in livestock farms. Nature Communications, 11(1):1427, 2020. doi: 10.1038/s41467-020-15222-y. URL https://doi.org/10.1038/s41467-020-15222-y.

[76] R. L. Tatusov, N. D. Fedorova, J. D. Jackson, A. R. Jacobs, B. Kiryutin, E. V. Koonin, D. M. Krylov, R. Mazumder, S. L. Mekhedov, A. N. Nikolskaya, B. S. Rao, S. Smirnov, A. V. Sverdlov, S. Vasudevan, Y. I. Wolf, J. J. Yin, and D. A. Natale. The COG database: an updated version includes eukaryotes. BMC Bioinformatics, 4(1):41, 2003. doi: 10.1186/1471-2105-4-41. URL https://doi.org/10.1186/1471-2105-4-41.

[77] E. A. Taylor, E. R. Jordan, J. A. Garcia, G. R. Hagevoort, K. N. Norman, S. D. Lawhon, J. M. Piñeiro, and H. M. Scott. Effects of two-dose ceftiofur treatment for metritis on the temporal dynamics of antimicrobial resistance among fecal escherichia coli in holstein-friesian dairy cows. PloS one, 14(7):e0220068–e0220068, 07 2019. doi: 10.1371/journal.pone.0220068. URL https://pubmed.ncbi.nlm.nih.gov/31329639.

[78] A. Tett, K. D. Huang, F. Asnicar, H. Fehlner-Peach, E. Pasolli, N. Karcher, F. Armanini, P. Manghi, K. Bonham, M. Zolfo, F. De Filippis, C. Magnabosco, R. Bonneau, J. Lusingu, J. Amuasi, K. Reinhard, T. Rattei, F. Boulund, L. Engstrand, A. Zink, M. C. Collado, D. R. Littman, D. Eibach, D. Ercolini, O. Rota-Stabelli, C. Huttenhower, F. Maixner, and N. Segata. The prevotella copri complex comprises four distinct clades underrepresented in westernized populations. Cell Host & Microbe, 26(5):666–679.e7, 2019. doi: https://doi.org/10.1016/j.chom.2019.08.018. URL https://www.sciencedirect.com/science/article/pii/S1931312819304275.

[79] C. Tong, D. Xiao, L. Xie, J. Yang, R. Zhao, J. Hao, Z. Huo, Z. Zeng, and W. Xiong. Swine manure facilitates the spread of antibiotic resistome including tigecycline-resistant tet(x) variants to farm workers and receiving environment. Science of The Total Environment, 808:152157, 2022. ISSN 0048-9697. doi: https://doi.org/10.1016/j.scitotenv.2021.152157. URL https://www.sciencedirect.com/science/article/pii/S0048969721072338.

[80] P. Trinh, D. S. Clausen, and A. D. Willis. happi: a hierarchical approach to pangenomics inference. bioRxiv e-prints, Apr. 2022. doi: https://doi.org/10.1101/2022.04.26.489591. URL https://github.com/statdivlab/happi.

[81] L. Van Gompel, R. E. Luiken, R. B. Hansen, P. Munk, M. Bouwknegt, L. Heres, G. D. Greve, P. Scherpenisse, B. G. Jongerius-Gortemaker, M. H. Tersteeg-Zijderveld, S. Garćıa-Cobos, W. Dohmen, A. Dorado-Garćıa, J. A. Wagenaar, B. A. Urlings, F. M. Aarestrup, D. J. Mevius, D. J. Heederik, H. Schmitt, A. Bossers, and L. A. Smit. Description and determinants of the faecal resistome and microbiome of farmers and slaughterhouse workers: A metagenome-wide cross-sectional study. Environment International, 143:105939, 2020. ISSN 0160-4120. doi: https://doi.org/10.1016/j.envint.2020.105939. URL https://www.sciencedirect.com/science/article/pii/S0160412020318948.

[82] A. K. Vidaver. Uses of Antimicrobials in Plant Agriculture. Clinical Infectious Diseases, 34 (Supplement 3):S107–S110, 06 2002. ISSN 1058-4838. doi: 10.1086/340247. URL https://doi.org/10.1086/340247.

[83] H. Wang, J.-S. Liu, S.-H. Peng, X.-Y. Deng, D.-M. Zhu, S. Javidiparsijani, G.-R. Wang, D.-Q. Li, L.-X. Li, Y.-C. Wang, and J.-M. Luo. Gut-lung crosstalk in pulmonary involvement with inflammatory bowel diseases. World journal of gastroenterology, 19(40):6794–6804, 10 2013. doi: 10.3748/wjg.v19.i40.6794. URL https://pubmed.ncbi.nlm.nih.gov/24187454.

[84] Y. Wang, N. Lyu, F. Liu, W. J. Liu, Y. Bi, Z. Zhang, S. Ma, J. Cao, X. Song, A. Wang, G. Zhang, Y. Hu, B. Zhu, and G. F. Gao. More diversified antibiotic resistance genes in chickens and workers of the live poultry markets. Environment International, 153:106534, 2021. doi: https://doi.org/10.1016/j.envint.2021.106534. URL https://www.sciencedirect.com/science/article/pii/S0160412021001598.

[85] H. M. Wexler. Bacteroides: the good, the bad, and the nitty-gritty. Clinical microbiology reviews, 20(4):593–621, 10 2007. doi: 10.1128/CMR.00008-07. URL https://pubmed.ncbi.nlm.nih.gov/17934076.

[86] A. Willis and J. Bunge. Estimating diversity via frequency ratios: Estimating diversity via ratios. Biometrics, 71, 06 2015. doi: 10.1111/biom.12332.

[87] A. Willis, J. Bunge, and T. Whitman. Improved detection of changes in species richness in high diversity microbial communities. Journal of the Royal Statistical Society. Series C (Applied Statistics), 66(5):963–977, 2022/06/26/ 2017. doi: https://www.jstor.org/stable/44682601. URL http://www.jstor.org/stable/44682601.

[88] D. E. Wood, J. Lu, and B. Langmead. Improved metagenomic analysis with kraken 2. Genome Biology, 20(1):257, 2019. doi: 10.1186/s13059-019-1891-0. URL https://doi.org/10.1186/s13059-019-1891-0.

[89] L. Wu, X. Xie, Y. Li, T. Liang, H. Zhong, J. Ma, L. Yang, J. Yang, L. Li, Y. Xi, H. Li, J. Zhang, X. Chen, Y. Ding, and Q. Wu. Metagenomics-based analysis of the age-related cumulative effect of antibiotic resistance genes in gut microbiota. Antibiotics, 10(8), 2021. ISSN 2079-6382. doi: 10.3390/antibiotics10081006. URL https://www.mdpi.com/2079-6382/10/8/1006.

[90] F. Xu, Y. Fu, T.-y. Sun, Z. Jiang, Z. Miao, M. Shuai, W. Gou, C.-w. Ling, J. Yang, J. Wang, Y.-m. Chen, and J.-S. Zheng. The interplay between host genetics and the gut microbiome reveals common and distinct microbiome features for complex human diseases. Microbiome, 8(1):145, 2020. doi: 10.1186/s40168-020-00923-9. URL https://doi.org/10.1186/s40168-020-00923-9.

[91] J.-Y. Yang, Y.-S. Lee, Y. Kim, S.-H. Lee, S. Ryu, S. Fukuda, K. Hase, C.-S. Yang, H. S. Lim, M.-S. Kim, H.-M. Kim, S.-H. Ahn, B.-E. Kwon, H.-J. Ko, and M.-N. Kweon. Gut commensal bacteroides acidifaciens prevents obesity and improves insulin sensitivity in mice. Mucosal Immunology, 10(1):104–116, 2017. doi: 10.1038/mi.2016.42. URL https://doi.org/10.1038/mi.2016.42.

[92] T. Yatsunenko, F. E. Rey, M. J. Manary, I. Trehan, M. G. Dominguez-Bello, M. Contreras, M. Magris, G. Hidalgo, R. N. Baldassano, A. P. Anokhin, A. C. Heath, B. Warner, J. Reeder, J. Kuczynski, J. G. Caporaso, C. A. Lozupone, C. Lauber, J. C. Clemente, D. Knights, R. Knight, and J. I. Gordon. Human gut microbiome viewed across age and geography. Nature, 486(7402):222–227, 2012. doi: 10.1038/nature11053. URL https://doi.org/10.1038/nature11053.

